# Disordered activation domains enhance DNA target search rate of a bZIP transcription factor

**DOI:** 10.1101/2025.11.15.688632

**Authors:** Jérémie Gaudez, Sarah L. Shammas

## Abstract

Transcription factors typically contain an effector domain and a DNA-binding domain, and are predominantly disordered. There is increasing evidence that effector/activation domains contribute to the DNA target search process. However, exactly how those domains influence this search is still unclear. Here, we have purified full-length CREB, a bZIP transcription factor with long disordered activation domains. We found, using *in-vitro* stopped-flow kinetic and equilibrium binding experiments, that the full-length protein has high dimerization affinity and dimerizes before binding DNA, as recently shown for the standalone CREB bZIP domain. CREB contains three recognised intrinsically disordered activation domains, which fall into classical acidic and glutamine-rich classifications. We show that all three contribute to decrease CREB’s general affinity for DNA. This, in turn, makes the target-search process more efficient by limiting CREB sequestration on non-target DNA. Thus, one of the ways in which intrinsically disordered, non-DNA-binding domains can affect target-search is by modulating non-specific DNA-binding affinity.

## Introduction

Transcription factors (TF) are proteins that regulate gene expression by binding to specific DNA motifs in the genome. Specific TFs typically contain at least two important domain types: the DNA binding domain (DBD) and the effector domain. The DBD adopts specific folds which are used to classify the factors into families, while effector domains regulate transcriptional activity, often by direct recruitment of coactivator partners and post-translational modifications (reviewed further in (1)).

Strikingly, TFs are often disordered or contain long disordered regions (1). Indeed, it is expected that around 90% of transcription factors contain large disordered regions (2), compared to 35% in the human proteome (3). Although disordered regions cannot adopt stable, long-lived structures independently under physiological conditions (4–6), they may achieve defined folds upon binding their partners, a phenomenon called folding upon binding (6–11). Disorder is proposed to offer many advantages for protein-protein interactions in general, such as multiple possible binding structures and partners (1,6), increased accessibility for binding (12) or post-translational modifications (13,14), and increased turnover (15) (meaning increased responsiveness to stimuli). Proposed mechanistic roles for protein disorder within transcriptional regulation are reviewed in detail in (16). Effector domains, which often expose Short Linear Motifs (SLiMs) that mediate protein interactions (17–19) and are generally acidic, are considered predominantly disordered and have been nicknamed “negative noodles” (1,20–22).

One important challenge transcription factors face is locating their limited target sites amongst the vast excess of non-specific DNA. It was hypothesised (23–27) and later observed (28–34), that search is optimised via facilitated diffusion. Facilitated diffusion consists of free 3D diffusion interspersed by sliding, hopping/jumping and intersegmental transfer. During target search, factors do seem to spend most of their time bound to DNA, suggesting non-target interactions dominate this process (35), yet *in vitro* studies tend to focus on target binding interactions.

Target search within the nucleus is largely DBD-driven for some transcription factors (36,37), but in others disordered non-DNA binding domains contribute to target search and DNA target binding (36–42). For example, domain swapping the IDRs of HIF-1α and HIF-2α concurrently changed their target-search behaviour, leading to distinct diffusive properties and showing a subset of DNA targets whose binding was dependent on the HIF isoform’s IDR and not their structured DBD (40). Compellingly, yeast transcription factor paralogs with different binding site preferences tended to have similar DBDs and differed instead in their non-DBDs (39). Yeast Msn2 disordered effector domains also independently localise to expected nuclear target sites, without DBDs (36), while the DBD alone of Msn2 (36) and other yeast factors (42) mostly bound to distinct sties compared to the wild-type factors. Effector domains are typically involved in protein-protein interactions, and it was recently proposed they direct a two-step target search, where they would direct factors by probing for the presence of partners near sites of interest (43). However, this appears insufficient for some factors, including Msn2, as its binding was seldom affected by the deletion of its Med15 coactivator or of other transcription-factors co-binding at overlapping promoters (although perhaps a key interacting partner is still missing) (42). Conversely, deletion of Msn2 and of its paralog Msn4 reduced the binding of multiple other transcription factors to Msn2/Msn4 promoters (42). Ultimately how disordered regions impact DNA target search is still unclear, and there are likely multiple mechanisms at play. Intramolecular interactions between acidic activation domains and DBDs could also play important auto-inhibitory roles in DNA binding and impact search.

CREB (cAMP response element binding-protein) is an essential transcription factor, whose deletion leads to perinatal death in mice and developmental defects (44,45). It is activated downstream of numerous signalling cascades by various kinases (46–53), most notably the cAMP cascade (54–56) via protein kinase A (PKA) which targets Serine 133 within CREB’s Kinase Activation Domain (KID) (46,49,54,57–59). This phosphorylation enhances binding to the KIX domain of CREB’s coactivator, CBP/p300 (CREB Binding Protein) thereby activating transcription (60). CREB also plays a role in memory retention (61–66) and in circadian rhythms (67). It is also overexpressed in many cancer types, indicative of tumour initiation, linked to worse prognosis, and responsible for tumour progression (57,68–70).

CREB binds to the palindromic CRE DNA sequence (TGACGTCA) and is a member of the bZIP transcription factor family. The bZIPs are named after their <u>b</u>asic <u>zip</u>per DNA binding and dimerization domain and are the second largest family of dimerising transcription factors (71). The bZIP is a bipartite domain, split into the DNA binding subdomain (basic region) and a dimerization domain (a helical leucine zipper) (72–76). bZIPs are typically disordered in solution but form helical coiled-coil domain structures as DNA-bound dimers (77–80).

CREB also has a large (∼260 residues) disordered transactivation domain (TAD) at its N-terminus (Figs. S1, S2). This domain is split into 3 subdomains: with glutamine-rich activation domains (Q1 and Q2) to either side of the acidic KID. Phosphorylated KID (pKID) forms a kinked helix when bound to CBP’s KIX (81) and is negatively charged, having many negatively charged residues flanking its hydrophobic binding motif (Fig. S3). The Q-rich domains have been largely neglected in biophysical and transcriptional studies. These disordered regions are known as constitutive activation domains, and their deletion decreases transcription from target promoters (82,83). Q2 appears critical for promoter occupancy and prolonged residence times (84), and binds the TAF4 subunit of TFIID (85,86). Both Q1 and Q2 are largely composed of uncharged, small residues (Fig. S3), and appear to contain small segments of transient β-structure (87), which may facilitate these interactions.

Unfortunately, very little is known about the individual impact of CREB’s corresponding domains on target search, besides a report that phosphorylation-dependent intramolecular interactions between the bZIP and pKID domains may decrease affinity for target DNA (88). There have also been no kinetic studies of the binding interactions of full-length CREB. Here, we present kinetic stopped-flow data that show CREB binding to its DNA target site remains extremely fast despite the presence of large, disordered flanking domains, and further uncover and rationalise a role for these effector domains in rapidity of target search.

## Results

### CREB readily forms homodimers and follows the dimer binding pathway

Previous studies have debated the dominant stoichiometry of CREB, suggesting it is largely monomeric (89) or largely dimeric (88,90) in solution; estimated equilibrium dissociation constants for homodimerization (*K*_D_,_dim_) vary over many orders of magnitude. Our recent work demonstrated that the isolated bZIP domain of CREB (see Fig. S1 for an overview CREB domain architecture) dimerizes before target binding at physiologically relevant concentrations, ostensibly because the CREB-bZIP monomer has a much higher affinity for itself than its target (91). Through altering kinetic rate constants for dimerization CREB’s activation domains might alter target search rates and stoichiometry of binding. Previously, kinetic rate constants for the isolated CREB-bZIP domain (henceforth referred to as simply bZIP) were determined using intrinsic fluorescence of a buried tyrosine as a probe (90,91). This strategy could not be extended to full-length CREB due to tryptophan residues elsewhere within the protein. Instead, fluorescently-labelled pseudo wild-type CREB was rapidly diluted 11-fold using stopped-flow apparatus to monitor relaxation kinetics to equilibrium, and thereby reveal the fundamental rate constants. A fluorophore was introduced via one of the naturally existing cysteine residues within the leucine zipper (C323), with both remaining cysteine mutated to serine which has no significant impact on DNA binding or dimerization (77,88). Upon dilution a significant increase in fluorescence intensity was observed, which depended upon the protein concentration in a manner expected according to a simple two-state model i.e. without populated intermediates. Simultaneous fitting of seven pairs of averaged kinetic traces (Fig. 1A, Fig. S4A-B) enabled simultaneous estimation of both homodimerization *K*_D_ (henceforth *K*_D,dim_) and homodimerization dissociation rate constant (henceforth *k*_2_) of (26 ± 6) nM and (0.10 ± 0.01) s^-1^ respectively (Fig. 1B). According to the two-state model applied the association rate constant (*k*_1_) was inferred to be (8 ± 2) µM^-1^s^-1^. Results were consistent for two different extrinsic fluorophores (AlexaFluor®488 and AlexaFluor®594) (Fig. S4C). Serial dilutions in SEC-MALS experiments confirmed CREB is fully dimeric in solution at µM concentrations, consistent with a *K*_D,dim_ in this nM range (Fig. S5A-B). These values are similar to those reported previously for the bZIP (88,90,91), and suggest that rapid collapse towards a dimer pathway might occur for the full length protein too.

**Figure 1:**
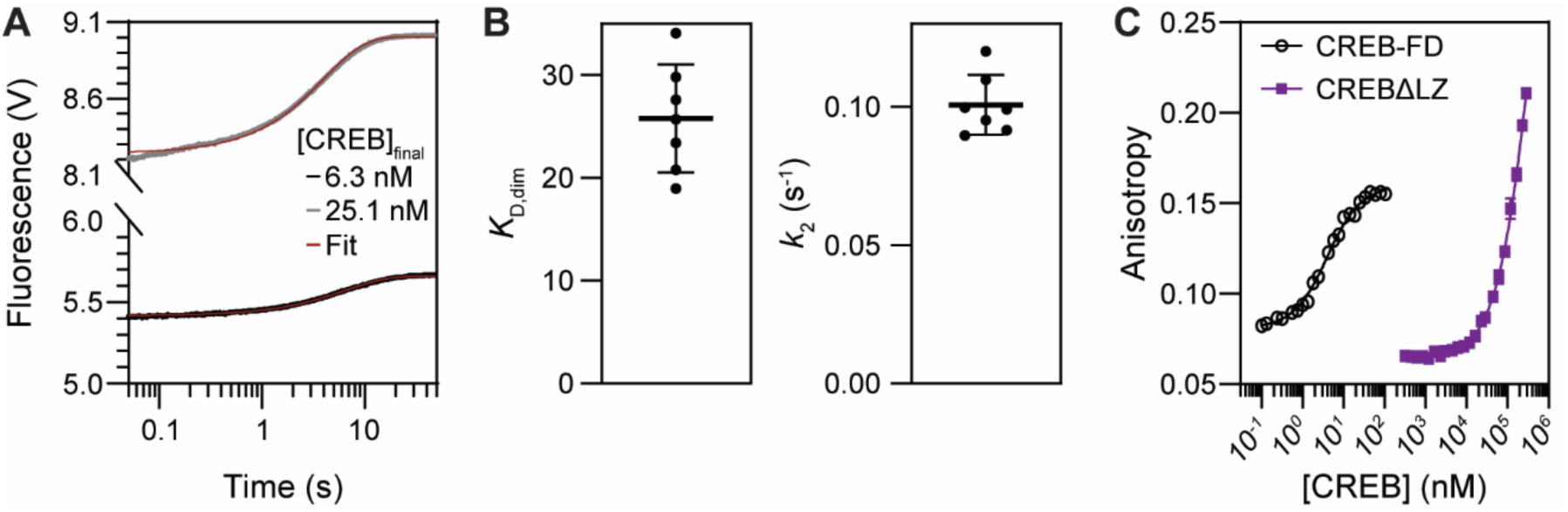
Monomeric CREB binds more tightly to CREB than its target-DNA CRE. **A.** Example pair of stopped-flow kinetic traces monitoring dilution-driven dissociation of AlexaFluor®488 labelled CREB homodimers. CREB solutions were 11-fold diluted into biophysical buffer to final concentrations of 6.3 nM (black, lower) and 25.1 nM (grey, upper). CREB complex dissociation was monitored by fluorescence intensity increase. Fits (brown lines) are to Eq. 5, where values of ΔF, *K*_D,dim_ and *k*_2_ are shared between the pair of datasets. All other fitted pairs, and fitting residuals, are shown in Fig. S4. **B.** Bar charts showing *K*_D,dim_ (left) and *k*_2_ (right) estimates for CREB for all pairs. Plotted error bars are standard deviation. **C.** Examples of fluorescence anisotropy equilibrium binding titrations measuring binding for CREB forced-dimer (CREB-FD, black open circles) and forced-monomer (CREBΔLZ, solid purple squares) to 1 and 5 nM AlexaFluor®488 labelled CREh respectively. Solid lines represent best fit to a single binding-site model (Eq. 3). Forced-dimer has high affinity for target CREh, displaying lower and higher plateaus, resulting in a reliable affinity estimate in the nM range. Replicate CREB-GGC curves are shown in Fig. S7. Anisotropy increase with CREBΔLZ only occurs at much higher concentrations, near the mM range, and may reflect increased solution viscosity rather than a binding event.

High homodimerization propensity does not guarantee that a protein follows the dimer pathway however. Flux through the two pathways at equilibrium is dictated by the ratio of *K*_D_,_dim_ to the equilibrium dissociation constant of monomer and DNA target (*K*_D, M-CRE_) (91). We designed forced-monomer mutants, by removing the leucine zipper (LZ) (Fig. S1). The resulting mutant, CREBΔLZ, was observed by SEC-MALS to be monomeric as anticipated (Fig. S5C). CRE binding affinity was examined by incubating AlexaFluor®488-labelled CRE hairpin oligos (AlexaFluor®488-CREh) with varying concentrations of CREBΔLZ. In such experiments fluorescence anisotropy typically increases as the DNA becomes proportionately more bound in complex, enabling an estimate of binding affinity. For the isolated bZIP the binding affinity is in the hundreds of µM range (91). However, fluorescence anisotropy was only increased at extremely high µM concentrations for CREBΔLZ, and did not plateau but continued to increase (Figs. 1C, S6A). Higher protein concentrations could not be examined due to protein aggregation concerns. The observed anisotropy increase may indicate very low affinity binding (*K*_D, M-CRE_ of around 1 mM) of multiple CREBΔLZ molecules, or increased solution viscosity due to the high protein concentrations. Additional methods were tested to attempt to discriminate between these possibilities. First, analogous fluorescence anisotropy equilibrium studies were performed where CREBΔLZ was fluorescently labelled, rather than the DNA. In this scenario the species present at high concentration is considerably smaller (one quarter the molecular weight), which should result in lower solution viscosity at any given concentration. The analogous equilibrium curve showed anisotropy increases only at even higher concentrations (Fig. S6A). ITC titrations with unlabelled CREBΔLZ and CRE DNA displayed no heat changes even for concentrations above 100 µM (Fig. S6B). Furthermore, although flow induced dispersion analysis (FIDA) experiments with AlexaFluor®488-labelled CREBΔLZ demonstrated increased apparent hydrodynamic radius upon incubation with mM concentrations of CRE DNA, these changes were fully explained by altered solution viscosity rather than DNA binding (Fig. S6C-D). This might suggest the observed fluorescence anisotropy increase with these same partners at mM DNA concentrations (Fig. S6A) was also due to viscosity changes rather than binding. A very cautious lower limit of *K*_D, M-CRE_ is therefore 1 mM (>10 mM base pairs). Monomeric full-length CREB binds extremely weakly to target DNA, yet retains similar dimerization kinetics to the isolated bZIP domain. Taken together, the results suggest full-length CREB also binds its target as a pre-formed dimer in DNA mixtures.

### CREB’s TAD decreases CREB’s binding rate with target DNA

Having established the importance of the dimer for target search, future effort focussed on the interaction of dimeric CREB with DNA. A covalent dimeric version of CREB (CREB-FD, Fig. S1) was generated by adding a GGC extension to the C-terminus of CREB and incubating under oxidising conditions. In contrast to monomer binding, dimeric CREB-FD binds tightly to AlexaFluor®488-CREh (*K*_D_ = (1.7 ± 0.8) nM) (Fig. 1C, Fig. S7, see Table S2 for oligo sequences). Binding resulted in increased fluorescence that could be used to probe reaction progress in stopped-flow mixing experiments (Fig. 2A). Kinetic traces were fit by a single exponential decay function (Fig. 2A), and observed association rate constants *k_obs_* depended linearly on protein concentration (Fig. 2B, Fig. S8). This behaviour is consistent with a simple two-state mechanism for dimeric CREB binding to target DNA, with the gradients in Fig. 2B representing the fundamental association rate constant, *k*_on_. CREB-FD has a *k*_on_ of (1.11 ± 0.04) nM^-1^s^-1^, roughly half that for a covalent dimeric version of the isolated bZIP domain (bZIP-FD) of (2.45 ± 0.08) nM^-1^s^-1^ (Fig. 2B-C, Table S1) under the same experimental conditions. Thus, the surrounding disordered activation domains act to slow target DNA binding. Interestingly the exact nature of the CRE-containing DNA sequence appears to have little effect on *k*_on_ for both CREB and bZIP, as mixing with a linear CRE-containing DNA construct rather than a hairpin has little to no effect on *k*_on_ (Fig. S9).

**Figure 2:**
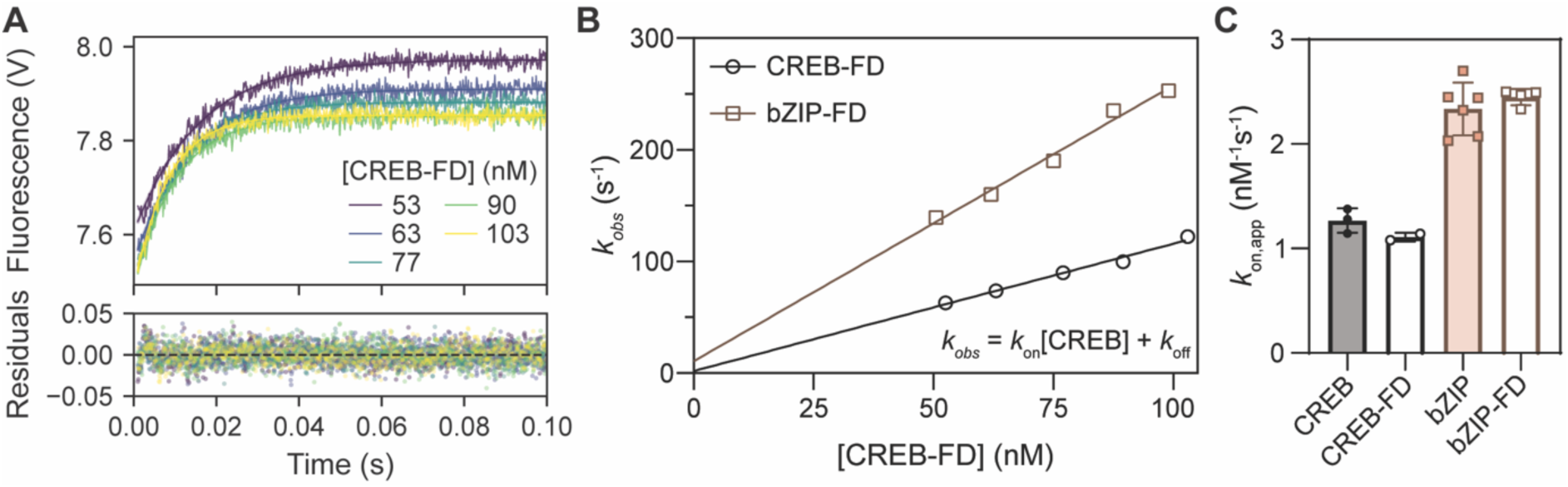
CREB’s bZIP has a higher association rate constant with target DNA than the full-length protein. **A.** Representative examples of association traces fit with a single-exponential decay function. Associated residuals from the fit are displayed beneath. **B.** Linear dependence of observed rate constant *k_obs_* on protein concentration. Lines represent the best straight-line fits to the data, where the gradient represents the association rate constant *k*_on_. **C.** Bar chart showing the *k*_on_ average for each mutant from the plotted repeats. Plotted error bars are standard deviation. *k*_on_ of the bZIP mutants are approximately twice that of CREB mutants. Forced-dimerisation appears to have negligeable effect on association kinetics indicating CREB is mostly dimeric prior to binding DNA.

Values obtained for non-forced-dimer pseudo wild-type CREB and bZIP (Fig. 2C, Table S1), are similar once corrected for dimer concentration, consistent with CREB being mostly dimeric prior to mixing with DNA. Without dimer concentration correction the y-intercept, which gives an approximation of *k*_off_, gives a slightly negative value. It is increased from a non-physical negative value towards 0 when higher CREB concentrations are used to reduce the relative monomeric population (Fig. S8). The y-intercept for CREB and bZIP is further dependent on the cutoff used in data analysis (Fig. S10; a 1 ms cutoff provides superior fit and yields a *k*_on_ close to that of CREB-FD (Fig. 2C)). These data further support the dimer pathway as sufficient to explain target binding *in vitro*, and also demonstrate an efficient method of calculating DNA binding parameters without the need for covalent dimerization.

### CREB’s activation domains assist target search

Although non-DNA binding domains impede association to target DNA in isolation, within the nucleus there are vast concentrations of competing non-specific DNA that complicate the process. To better mimic the process of CREB target search for CREB, we performed analogous binding experiments to those described above, but in the presence of large excesses of unlabelled non-specific dsDNA (nCRE). To allow maximal control and develop an accurate quantitative understanding of how non-specific DNA interferes with binding, we used a single non-specific DNA oligo that contained no half or full CRE sites. Experiments were performed at multiple [nCRE] up to 10 µM, which is equivalent to 140 µM base pairs. As previously, kinetic traces were fit by single exponential decay functions (Fig. S11A), and extracted rate constants depended linearly on protein concentration (Fig. 3A, Fig. S11B-C). For both dimeric bZIP-FD and CREB-FD target binding rates are reduced by the inclusion of competing non-specific DNA. Remarkably, although bZIP-FD has higher *k*_on_ than CREB-FD in the absence of competitor, apparent search rates drop more steeply with competitor concentration for bZIP-FD than for CREB-FD (Fig. 3B). At the highest competing DNA concentrations, target binding rates for CREB-FD are similar, or even higher than those for bZIP-FD.

**Figure 3:**
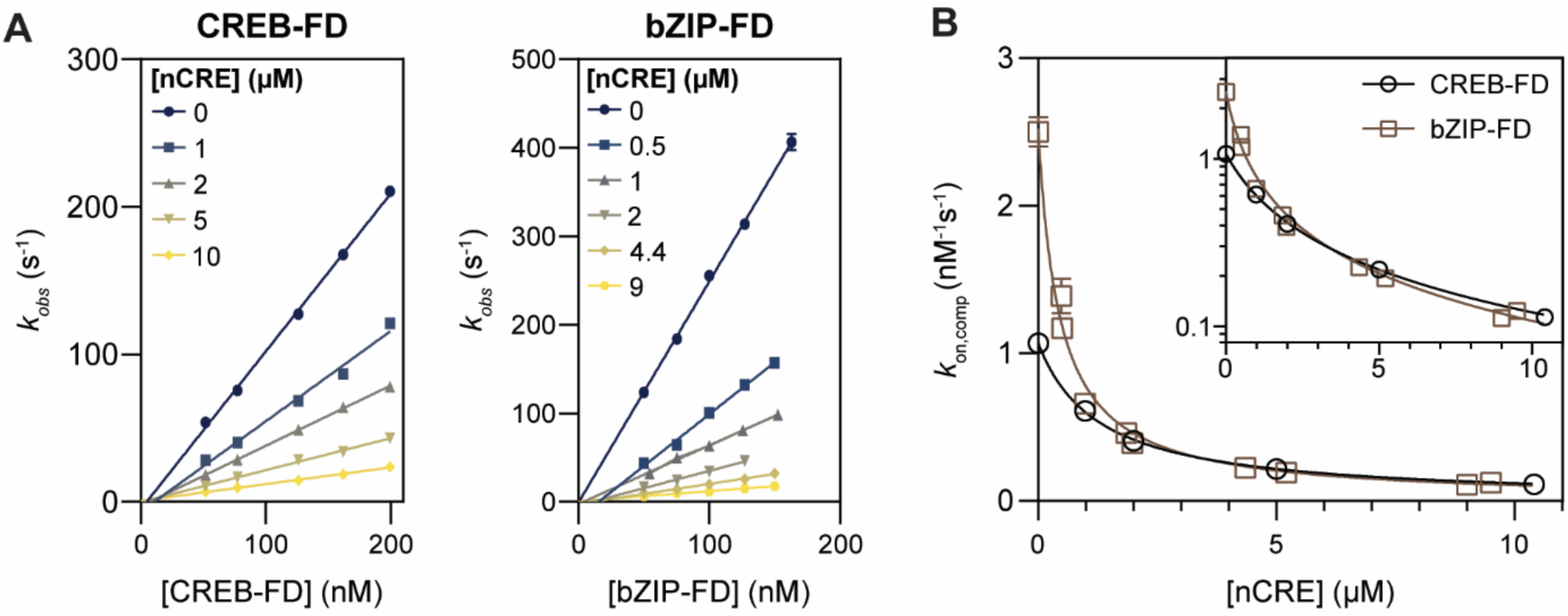
Competing non-specific DNA slows target binding by CREB. **A.** Linear dependence of observed rate constants *k_obs_* on protein concentration in association experiments for CREB-FD (left) and bZIP-FD (right) with 5 nM AlexaFluor®488-labelled CREh. Lines represent best straight-line fits to the data, where the gradient represents *k*_on,comp_. Concentrations of competing non-specific DNA (nCRE) are indicated by colour according to the inset legend. All concentrations are those within the observation cell. **B.** Decrease of apparent association rate constant, *k*_on,comp_, with increasing competitor concentration. Lines represent the best fit to a model where CREB is sequestered by competitor DNA (Eq. 4). Inset shows the same data on a log-scale.

We reasoned that this behaviour could be rationalised by a difference in the concentration of CREB available to bind DNA. At high competitor concentrations a significant proportion of CREB may be sequestered on competitor, leaving only a limited proportion to bind target DNA. Indeed, a simple model using the *K*_D_ of CREB binding to nCRE (*K*_D,nCRE_) is able to model the observed behaviour quite well in both cases (Fig. 3B). The *K*_D,nCRE_ for CREB-FD and bZIP-FD extracted from the fits are (1.21 ± 0.02) µM and (0.45 ± 0.04) µM respectively. These indirect estimates from modelling are consistent with independent estimates from ITC of (1.3 ± 0.4) µM and (0.36 ± 0.03) µM respectively (Fig. S12, Table S1). In other words, target search rate constants in the different DNA mixtures can be predicted using this model and ITC estimates of *K*_D,nCRE_ (Fig. S11F). Plotting *k*_on,comp_ / *k*_on_ against predicted [CREB]_free_/[CREB]_total_ calculated using ITC estimates of *K*_D,nCRE_ reveals a straight line, also demonstrating that binding rates reflect the free CREB concentration (Fig. S11D). As previously, bZIP and CREB display the same behaviour as their forced dimeric counterparts, indicating the presence of the competitor has not significantly changed the binding stoichiometry (Fig. S11). The model might be further used to infer the behaviour at even higher [nCRE]; for context the highest experimental DNA concentration is roughly 100-fold lower than the base pair concentration within the Eukaryotic nucleus. In the presence of such high competitor concentration we would expect binding by the full-length protein to be more rapid than the bZIP. Hence, it appears that the activation domains ameliorate the retardation of DNA target binding by competing DNA, by decreasing CREB’s affinity for non-target DNA.

### Activation domains collaborate to decrease DNA binding affinity and rates

A series of domain deletion mutants were generated to uncover the domain (or domains) responsible for ameliorating the retardation of target binding rates in DNA mixtures. CREB with Q1, KID, and Q2 deletions, referred to as CREBΔQ1, CREBΔKID and CREBΔQ2 were expressed and purified for characterisation of DNA binding parameters. Retardation is controlled through *K*_d,nCRE_. Unfortunately, the CREBΔKID mutant was particularly prone to aggregation at high concentrations so *K*_d,nCRE_ could not be characterised by ITC. An alternative mutant was generated where the KID domain was replaced by an equivalent-length GS-linker to assess any sequence-specific effects (CREB-GS-KID, Fig. S1). The effects of all mutations on *K*_D,nCRE_ are sizeable, being 2-3-fold lowered (Figs 4A, S12). Thus, it appears that all activation domains contribute to the ameliorated retardation. The effects of mutation appear broadly additive, as seen by the *ΔΔG* values (*ΔG*_CREB_ - *ΔG*_mutant_ calculated from the ITC *K*_D,nCRE_ values) (Fig. S13).

**Figure 4:**
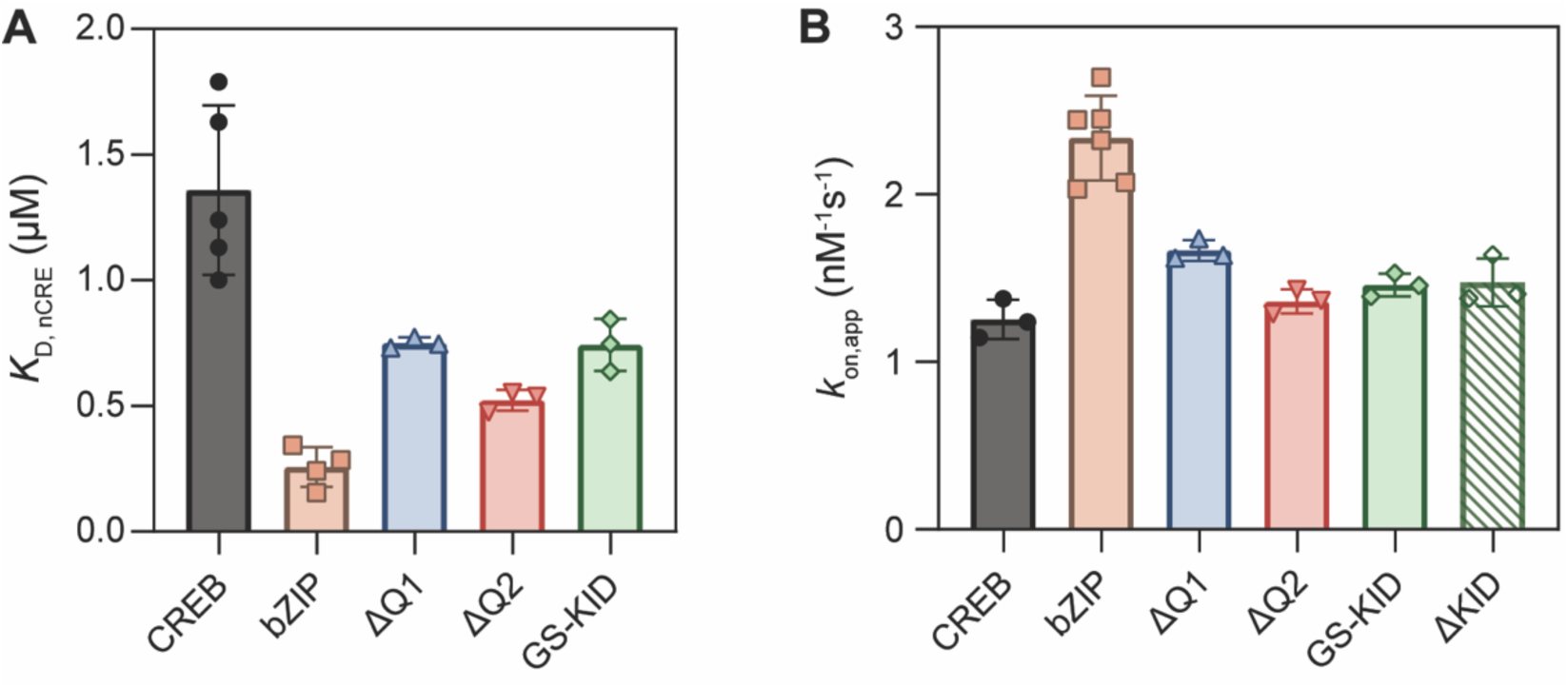
CREB’s activation domain all contribute to decrease DNA binding. **A**. Bar chart of *K*_D,nCRE_ estimated from replicate ITC experiments. Individual ITC experiments are shown in Fig. S12. Error bars are standard deviation in all panels. No individual domain mutation recapitulates the bZIP behaviour for either parameter. **B**. Bar chart showing average *k*_on_ from replicate concentration-dependent association experiments. Data points are calculated as gradients of straight-line fits shown in Fig. S8. *k*_on_ was determined assuming all mutants follow the dimer pathway, so are for dimeric concentrations.

Target search rates are determined by a combination of *k*_on_ and *K*_D,nCRE_ so target association rate constants were also assessed. All domain deletions slightly increased *k*_on_ for labelled CREh (Fig. 4B), but not to the level achieved by the isolated bZIP domain. Truncation reduces the size of the protein, so some increase in diffusional coefficient is anticipated, which would increase *k*_on_ correspondingly. Indeed, there is an apparent correlation between these parameters (p = 0.01, n = 6), though CREB-GS-KID and CREBΔKID have very similar *k*_on_ (Fig. 4B, Fig. S14). Truncation also significantly affects the net charge of the protein, and there is a correlation between these two parameters as well (p = 0.01, n = 6) (Fig. S14).

## Discussion

### Role of activation domains in DNA binding stoichiometry

The results of this study add to mounting evidence (88,90,91) against claims that CREB is monomeric in solution and cooperatively assembles on target DNA (89). Recently we showed that the combination binding rate constants for CREB bZIP favour a very rapid convergence to an almost entirely dimeric mode of target complex formation (91). As a eukaryotic transcription factor, CREB contains long disordered activation domains however. Post-translational modifications to these activation domains, aside from altering interactions with protein partners such as coactivators, can provide a mechanism to alter DNA binding affinities (92), but also potentially their target search stoichiometry. Such interference might be particularly expected from classical negatively-charged activation domains, such as KID, whose presence could impact homodimerisation of the positively charged bZIP domain. Consistent with affinity estimates from mass photometry (88), we find the inclusion of the activation domains to make little difference to equilibrium homodimerisation properties. However, the association rate constant for dimer formation (*k*_1_) will play an important role in the rapidity with which the dimer pathway is converged upon. Association rate constants for protein-protein interactions vary over many orders of magnitude; that for the isolated bZIP is on the higher end of typical (91,93). Should flanking activation domains reduce the inherent dimerisation rate constants then this would leave space for high monomer pathway flux within a kinetically-controlled system. However, the results here have shown the disordered activation domains of CREB also have little influence on dimerization kinetics, and therefore the rate at which equilibrium is approached, at least *in vitro*. Instead, the TAD appears to further decrease the affinity of the monomer for the DNA target to the extent we have been unable to quantify it. This effect clearly disfavours further the efficiency of the monomer pathway. Recently our lab showed that isolated bZIP domains may share similar kinetic and equilibrium parameters, and therefore may also follow a dimer pathway *in vitro* (91) (unlike previous reports (73,89,94–96)). However CREB is one of the few bZIPs to almost exclusively form homodimers (97), and is highly expressed in the cell as a baseline (98). Many bZIPs form heterodimers (e.g. Jun and Fos) or homodimers with lower affinity, and are present at lower levels in cells (97,98). This may relate to a difference in pathway preference, and it remains possible that this could vary by bZIP transcription factor, with control being exerted through the activation domains. Further experiments with other full-length proteins would be needed to establish this.

### Role of activation domains in rapidity of DNA binding

A significant decrease in association rate constant in the context of the full-length protein is expected. Most obviously, the protein will diffuse more slowly in solution due to its increased size, thereby decreasing the encounter rate with DNA. Hydrodynamic radii (R) of disordered proteins have a more complex dependence upon the number of residues (N) than those of folded proteins, and vary greatly depending upon sequence composition and experimental conditions e.g. ionic strength and temperature. If *R* = *N*^𝛾^, then 𝛾 typically ranges between 1/3 and 3/5 (99). Thus, R might be 1.7-2.5-fold increased for CREB compared with bZIP, which would translate to an approximate 1.07-1.23-fold increase in encounter rate (if charges are neglected). These calculations are not expected to be accurate, but are presented to demonstrate the effect of size on truncation is expected to be smaller than the 1.8-fold observed. The DNA binding region of the protein constitutes a much smaller fraction of the protein in the full-length protein, which could reduce the success of binding upon encounter.

However, the net charge also correlates with *k*_on_, and this might be a better explanation for the observed differences, particularly since equivalently charged GS-KID and ΔKID have very similar *k*_on_. As observed previously for the isolated bZIP (91), the association rate constants are extremely high, in the range typically referred to as diffusion-limited, and indicative of electrostatic rate enhancement. The charge of the full CREB molecule is -7 at neutral pH, quite different to that of the isolated bZIP domain at +7. That *k*_on_ remains so high for CREB despite its overall negative charge highlights the importance of clusters of charged residues along binding surfaces for electrostatic steering, rather than just net protein charge, in determining association rates. A dependence of *k*_on_ upon charge could indicate specific charge-mediated intramolecular interactions between domains. Previous NMR data have indicated interactions between a negatively-charged region of the TAD and the positively-charged bZIP (92). Pre-existing intramolecular interactions might significantly interfere with favourable long-range electrostatic interactions driving the association between the bZIP and DNA, thereby decreasing the extent of electrostatic rate enhancement. This does not appear to be the mechanistic explanation of the differences however, since removing the KID domain entirely or swapping it for a neutrally charged GS-linker have only a minor impact on *k*_on_. It is possible these intramolecular interactions in CREB are insufficiently populated (at physiologically relevant ionic strengths) to significantly impact association rates; NMR also suggests such interactions are minimal in the absence of CREB phosphorylation (92). The data appear more consistent with overall changes in encounter rate due to perturbed long-range electrostatic forces.

### Role of activation domains in target search

Importantly, CREB’s non-DNA binding activation domains decrease non-specific DNA binding affinity, with a concomitant improvement in target search inside highly concentrated DNA mixtures. This could translate to a more efficient target-search in the nucleus, and provide a simple mechanism for flanking disordered domains to affect target search (36,40,42). Post-translational modifications within the activation domain could harness this behaviour and modulate affinity of DNA, potentially helping or hindering gene expression to certain targets. Previously it has been shown that acetylation of MITF transcription factor activation domains control activity and search kinetics by lowering overall DNA binding affinity (100). If disordered transactivation domains help accelerate target search by decreasing DNA binding affinity, this could represent a general importance in target search (36–40) in a manner that is independent of partner recognition and recruitment (42).

The KID has many negative residues, as is typical for activation domains (12,101–103), so is an obvious candidate for generating self-inhibition by competing for binding the positively charged DNA binding domain. Indeed, association of HMGB1 and Antp homeodomain to target DNA was accelerated in the presence of competitor by using negatively charged protein extensions (104). Such an auto-inhibition mechanism in DNA binding has been previously proposed for CREB (92). The activation domain contains a casein-kinase cassette which when increasingly phosphorylated, results in successively decreased affinity for its DNA target (92). Intramolecular interactions of this nature have been inferred from NMR measurements (87,92) and are observable in AlphaFold (105) structures for CREB (Fig. S2). Should this be the major determinant however, we might expect deletion of the KID domain to explain all, or the vast majority, of the difference in DNA binding affinity between CREB and the isolated bZIP. Instead, all of CREB’s non-DBDs combine to decrease binding affinity. Replacing the entire negatively charged KID domain with a neutral GS-linker increases DNA binding affinity by less than 2-fold, only around 1/3 of the observed effect. Since phosphorylation in the activation domain is reported to increase its interactions with the bZIP (92), it is possible that phosphorylated versions might show a significantly larger impact. Surprisingly deleting the glutamine-rich Q1 domain, which is neither adjacent to the DNA-binding bZIP domain nor complementary in its charge, has a similar impact. The largest change is generated by removing the Q2 domain between the KID and bZIP domains. This is especially relevant since the sequence of this protein resembles that of a natural isoform of CREB present in human testis (htCREB) that lacks the Q2 domain (106). Surprisingly htCREB was previously reported to bind as tightly to half-CRE sites as CREB does, so it is possible that specificity is increased (92). Significantly disrupted intramolecular interactions between KID and bZIP domains resulting from Q2 removal could explain some reduction in DNA binding affinity. However, this should make the affinity changes highly non-additive by “double counting” the removal of a bZIP-KID interaction; whereas free energy changes are additive within error. Affinity measurements with htCREB have also displayed higher levels of CK phosphorylation-dependent auto-inhibition (92), indicating deleting Q2 actually encourages bZIP-KID interactions rather than disrupts them. These observations do not support reduced bZIP-KID interaction as responsible for the increased DNA binding affinity upon deleting Q2. Thus, it appears Q1 and Q2 may make significant contributions to decreasing non-specific DNA binding affinity of their own accord. Thus, the entire TAD may promote delocalisation, decreasing promoter occupancy. This may contribute to explain previous observations that CREB TAD deletions of increasing length tend to correspondingly decrease basal transcription (82).

Specific effects on association rate constants with competitor DNA appear unlikely in the absence of those with target DNA (*k*_on_). Thus, since *k*_on_ is less sensitive to truncation than *K*_D,nCRE_, it is reasonable to infer that the changes in *K*_D,nCRE_ are also mediated through altered dissociation rates from competitor DNA i.e. the presence of the activation domains lowers the kinetic stability of the complex. At this point the mechanism remains unclear. The Q-rich domains of CREB have been recently demonstrated to interact, potentially via. transient beta-strands (87), but it is unclear why this would impact DNA residency time. Structural or dynamical changes may be induced inside the DNA binding domain. This might be supported by the largest impact being the deletion of the neighbouring Q2 domain. Alternatively, or in addition, intramolecular interactions for CREB may compete and actively promote dissociation; an effect that might be further modulated by post-translational modifications. Given that the activation domains fall into two separate classes in terms of sequence composition it is possible that more than one mechanism is acting.

## Concluding remarks

Activation domains are required in addition to DNA binding domains for efficient transcription. Their incorporation into transcription factors can slow target binding. However, we show here that this apparent penalty can disappear within DNA mixtures if they simultaneously reduce sequestration on non-specific DNA. If anything, full-length transcription factor could bind targets somewhat faster than smaller isolated DNA binding domains at very high DNA concentrations, such as those within the nucleus. Activation domains have recently come under scrutiny for modulating cognate DNA binding sites and target search processes within nuclei (36–42). Our finding reinforces the notion that non-DNA binding domains can affect target search, even in simple DNA mixtures.

## Methods

### Cloning

A synthetic gene for a modified version of CREB isoform B (Uniprot P16220-2, basis for amino acid numbering) was fused with a DNA sequence encoding an N-terminal 6xHis Tag-containing SUMO solubility tag, and cloned into a pET vector (107,108). The modification involved mutation of residues C286, C296 and C323 to serine to improve solubility without affecting DNA binding properties (77,91). A gene for CREB-FD was subsequently generated from this gene by introducing a DNA sequence encoding a C-terminal GGC extension, which allows for covalent dimerization by disulfide bond formation under oxidising conditions. The vector was also used as a basis to generate vectors containing sequences for the CREBΔQ2, CREBΔKID, CREBΔLZ, and CREB-GS-KID mutants (Fig. S1).

Cysteines were reintroduced at their endogenous positions (C323 for full-length CREB, C296 for CREBΔLZ) using Q5 site-directed mutagenesis (New England Biolabs) with primers ordered from Life Technologies. The bZIP domain (residues 263-325) and bZIP-FD were expressed from pre-existing vectors, which encode CREB sequences fused with an N-terminal 6xHis Tag-tagged B1 domain of Streptococcal protein G (GB1), cloned into a pRSET-A vector (91). A sequence encoding the tobacco etch virus (TEV) protease site is situated between CREB and GB1 encoding sequences. This base vector was used to generate a vector encoding the CREBΔQ1 mutant (Fig. S1). Mutations were either generated by Q5 site-directed mutagenesis using primer ordered from Life Technologies, or NEBuiler® HiFi assembly (New England Biolabs) following manufacturer’s instructions and using inserts ordered from Integrated DNA Technologies) and verified by DNA sequencing. Protein charge for the clones was calculated as the difference between positive and negatively charged residues at neutral pH given by the ProtParam webserver (109).

### Protein expression

For expression of CREB and its mutants, plasmids were transformed into chemically competent *Escherichia coli* Rosetta2 (DE3) and selected by antibiotic resistance. Cells were grown into auto-induction 2xTY medium at concentrations of 0.5% v/v glycerol, 0.05% w/v glucose and 0.2% w/v lactose. Cells were grown at 30 °C, 180 rpm until OD_600_ of 0.6 was reached, when temperature was lowered to 18 °C for overnight recombinant protein expression. For expression of bZIP, bZIP-FD and CREBΔQ1, plasmids were transformed into *Escherichia coli* BL21(DE3)pLysS. Cells were cultured in 2xYT medium supplemented with 0.2% (w/v) glucose until the optical density (OD_600_) reached 0.6-0.8, expression induced with 1 mM IPTG, and the culture incubated for another 4 h at 37 °C.

### Protein purification

bZIP and bZIP-FD mutants were purified as previously described (91). For purification of CREB and mutants, each 4g pellet of cells harvested by centrifugation was lysed by sonication in 50 ml of buffer A (100 mM Tris pH 8.5, 50 mM NaCl, 1 mM DTT, 1x Pierce EDTA-free protease inhibitors (Thermo Scientific)). Lysate was heat-treated at 75 °C for 30 minutes before resting on ice for 10 minutes. Precipitate was cleared by centrifugation and the supernatant loaded on a Cytiva 5 mL pre-packed HisTrap™ HP columns pre-washed with buffer A. The column was then consecutively washed with 40-column volumes (CV) buffer A, 60 CVs of buffer B (100 mM Tris pH 8.5, 2 M NaCl, 1 mM DTT), and 5-CVs of buffer A again. Protein was eluted with a linear gradient of 0-1.5 M imidazole in buffer A. Fractions containing CREB were pooled, and TEV (CREBΔQ1 only) or ULP-1 protease added, then immediately dialysed into buffer C (10 mM sodium phosphate pH 8, 2 mM DTT) supplied with 1 mM EDTA. ULP-1 cleavage leaves an N-terminal scar with sequence GAMGS, and TEV cleavage a SDNAIA scar. The protein was loaded on a Cytiva HiTrap™ heparin 5mL pre-packed column equilibrated with buffer C. Flowthrough was reloaded on the column once to ensure complete binding of the protein to the column, before a 5 CV wash with buffer D (10 mM sodium phosphate pH 8, 2 mM DTT, 8 M Urea) followed by elution against a 0-500 mM NaCl gradient in buffer D. Pooled fractions were analysed by SDS-PAGE. If contaminants were still present, eluate was dialysed into buffer E (20 mM Tris pH 7.6, 2 mM DTT) and loaded on a Cytiva HiTrap™ Q HP 5 mL pre-packed column, equilibrated with buffer E. Protein was eluted using a 0-2 M NaCl gradient in buffer E. bZIP and bZIP-FD and purification were carried out as previously described (91). Purity was assessed after every chromatography step by SDS-PAGE and A260:A280 absorbance ratio.

### Generation of covalent dimeric CREB

To oxidise the GGC extension to form dimers, proteins were concentrated to 50 µM or higher, mixed with an equal volume of 100 mM Tris-HCl buffer (pH 9.2), placed in the tubes with loose fitting caps and kept in a shaking incubator for 16 h at 37 °C. CREB-FD underwent an extra denaturing anion-exchange chromatography step to separate covalent dimers from monomers. This was identical to the procedure described above for purification of CREB, but with all buffers supplemented with 8 M urea. GGC forced dimerization was confirmed both by SDS-PAGE and by LC-MS mass spectrometry.

### Protein labelling

CREB was fluorescently labelled using maleimide chemistry via. C323 for CREB (equivalently C296 for CREBΔLZ). CREB was diluted to 10-20 µM and dialysed in labelling buffer (20 mM Tris pH 7.5, 500 mM NaCl) supplemented with 1 mM TCEP for reduction of cysteines. TCEP was removed by buffer exchange into labelling buffer using a Cytiva PD-10 gravity column. Protein was then incubated with a 4-fold molar excess fluorophore, either AlexaFluor®488 maleimide or AlexaFluor®594 C5 maleimide, overnight for 4°C on a centrifuge tube roller. Reaction was quenched with 1 mM DTT. Dye, unlabelled and labelled proteins were dialysed post-labelling into buffer E and separated using anion-exchange chromatography, as described above for CREB.

### Preparation of CREB solutions for biophysics

Purified protein was twice dialysed against 1 L biophysical buffer (10 mM HEPES pH 7.4, 150 mM NaCl, 10 mM MgCl_2_, 0.05% (v/v) Tween-20). Protein concentration was recorded through UV-Vis spectroscopy before flash-freezing in liquid nitrogen and storage at -80°C. Extinction coefficients were determined, using the Gill, Von Hippel method (110) to be, for monomeric constructs: CREB – 9000 ± 195 M^-1^s^-1^, bZIP – 2630 ± 26 M^-1^s^-1^, CREBΔQ1 – 8200 ± 150 M^-1^s^_1^, CREBΔQ2 – 3500 ± 20 M^-1^s^_1^, CREBΔLZ – 8000 ± 250 M^-1^s^-1^, CREB-GS-KID – 8120 M^-1^s^-1^; and for dimeric constructs: CREB-FD – 19900 ± 400 M^-1^s^-1^, bZIP-FD – 5450 ± 100 M^-1^s^-1^. Accurate CREBΔKID concentrations were calculated by diluting into denaturant to avoid aggregation at concentrations required for UV-vis spectroscopy. For accuracy, all dilutions were performed gravimetrically.

### Preparation of oligonucleotides for biophysics

Oligonucleotides were purchased (Thermo Fisher or Integrated DNA Technologies). Oligonucleotide sequences are listed in Table S2. DNA was resuspended in annealing buffer (30 mM HEPES pH 7.5, 100 mM potassium acetate) at 100 µM. Oligos were annealed by denaturation for 5 minutes at 92 °C before slowly ramping down incubation temperature by 0.1 °C/s until 4 °C. Removal of unannealed or incomplete oligos was carried out by ion-exchange chromatography. Annealed products were diluted in buffer F (50 mM Tris pH 7.2, 1 mM EDTA), loaded on a HiTrap Q HP 5 mL pre-packed column pre-equilibrated with buffer F and eluted with a 0-1.5 M gradient NaCl in buffer F. Oligos were concentrated and buffer exchanged into biophysical buffer using Amicon centrifugal filter units before storage at -20°C.

### Data analysis

Curve fitting was performed on Python3 using the lmfit package (111) or on GraphPad Prism (version 10.0.0 for Windows, www.graphpad.com), using the Levenberg-Marquardt least-squares algorithm.

### Fluorescence anisotropy equilibrium binding titrations

Samples were incubated for at least 1 hour, protected from light, and then transferred to a 3 mm pathlength quartz cuvette (Hellma^®^). Fluorescence measurements were made using a Horiba FluoroMax-4 Spectrophotometer with a fluorescence polarization accessory, maintained at 25.0 °C. Excitation/emission wavelengths were 495/519 nm for AlexaFluor®488, and 590/618 nm for AlexaFluor®594. Three to ten fluorescence anisotropy measurements were averaged for each sample. Intensity, if not recorded directly, was calculated using the following equation (112):

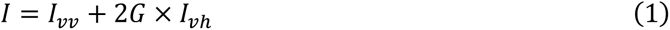

Where 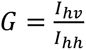 is the G factor, and *I*_xx_ are polarised intensities given by the fluorimeter. If an intensity change was noticed between bound and unbound species, anisotropy and intensity were fit by a single-site model and anisotropy corrected following the formula (112,113) (Equation 2):

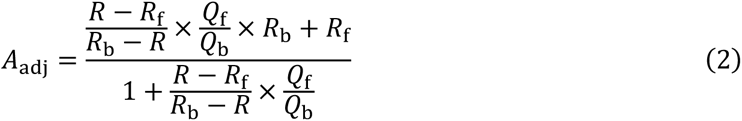

Where *R* is the anisotropy measurement of the sample, Q is the intensity measurement of the sample, and the subscripts *f* and *b* denote values measured for the free and bound sample, respectively.

Anisotropy binding titrations were fit to the following equation, applicable for a standard two-state binding model for a single site, as per (112,114):

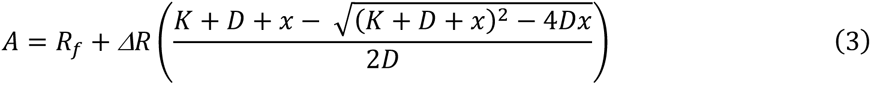

where *A* is the measured anisotropy*, R_f_* is the the initial anisotropy value, *ΔR* is the difference between the saturated and initial anisotropy value, *K* is the *K*_D_ in nM of DNA binding, *D* is the labelled probe concentration (typically DNA) in nM, and *x* is the titrant concentration (typically CREB), in nM.

### Stopped-flow kinetics

Kinetics measurements were performed on an Applied Photophysics SX-20 fluorescence stopped-flow spectrometer, at 25 ± 0.1 °C. Excitation wavelengths were 495 nm for AlexaFluor®488, and 590 nm for AlexaFluor®594. Slit widths were 2 mm for both excitation and emission. Emitted light was filtered by long-pass filters; 515 nm for AlexaFluor®488 and 610 nm for AlexaFluor®594. Appropriate timescales for traces were chosen to be approximately 10 half-lives (112).

Association rate constants (k_on_). CREB (200-800 nM) was rapidly mixed in a 1:1 volume ration with 10 nM AlexaFluor®488-CREh containing 0-20 µM nCRE (1-20 µM). A pressure-hold was applied to the syringes, and the first 1-3 ms of data discarded to account for mixing time. 50+ individual traces were averaged, and fit to a single exponential decay function to extract observed rate constants (*k_obs_)*. The apparent association rate constant, was calculated as the gradient of a straight line fit of *k*_obs_ against CREB concentration. When nCRE is absent this rate constant is equal to the fundamental association rate constant, *k*_on_. In the presence of nCRE, this rate constant is termed *k*_on,comp_. It is decreased from *k*_on_ due to sequestration of CREB on nCRE DNA according to Equation 4, which is derived in the Supplementary Materials:

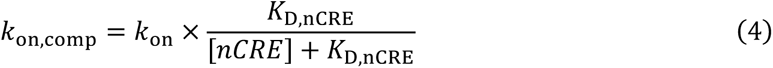

CREB homodimer dissociation (*k*_2_, *K*_D,dim_). 10.8-200 nM fluorescently-labelled CREB was rapidly diluted with biophysical buffer in a 10:1 volume ratio to encourage homodimer dissociation. Due to required timescales of data collection a pressure-hold could not be applied to the syringes, and therefore the first 100 ms of data was rejected to avoid mixing artefacts. 30+ kinetic traces were averaged. Averaged traces from solutions with different CREB concentrations were paired. Paired datasets were collected on the same day, without changing instrument calibration. Paired kinetic traces were simultaneously fit to Equation 5 (91,115):

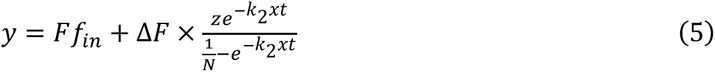

where:

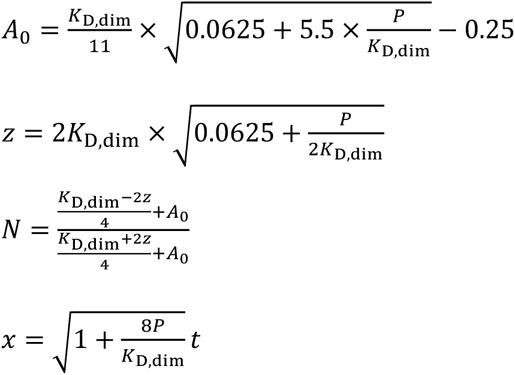

and *K*_D,dim_ *is* the homodimerization dissociation constant, *P* is final protein concentration, *y* is the fluorescence intensity, *t* is the time since mixing, and *k*_2_ is the dissociation rate constant of CREB homodimer. ΔF is the fluorescence intensity change and *Ff_in_* is the fluorescence signal at the end of the experiment.

### Isothermal titration calorimetry (ITC)

Protein and DNA were dialysed overnight in biophysical buffer together. ITC titrations were performed on a Malvern Panalytical MicroCal PEAQ-ITC with 13 to 19 injections. Protein was placed in the reaction cell and DNA in the syringe. Cell protein concentration was chosen to have a c-value (the product of the stoichiometry parameter and macromolecule concentration, divided by the dissociation constant) greater than 10. This was achieved by using bZIP and bZIP-FD starting dimeric concentrations of 7.5 to 21 µM and nCRE concentrations of 57-200 µM, and starting dimeric protein concentrations of 10-25 µM and starting nCRE concentrations of 126-206 µM. In one exceptional case, a low c-value titration was performed with 2 µM CREB-FD and 21 µM nCRE starting concentrations (Fig S12, run 3).

The following conditions were kept across all runs: 25.0 °C with high feedback, 750 rpm stir speed, 60 seconds delay, 150 seconds injection spacing, 4 seconds injection duration. Each titration was control corrected by dilution of the same titrant solution into biophysical buffer. Titrations were plotted and fit to a single-site binding model using in-built equations from the MicroCal PEAQ-ITC analysis software.

## Acknowledgements

UK Medical Research Council [MR/N024168/1 to S.L.S.];

