## Supplementary Materials for "Disordered activation domains enhance DNA target search rate of a bZIP transcription factor"

**Table S1:** Association rate constants to Alexa Fluor® 488-labelled CREh and dissociation constants of binding to unlabeled nCRE of the CREB constructs used in this study. All values are averages from at least triplicate with standard deviation error, except signaled with * where it is the average of 2 replicates, with error the difference between the average and the individual replicates.

| Construct | $k_{on}$ ($nM^{-1}s^{-1}$) | $K_{D,nCRE}$ from ITC (µM) | $K_{D,nCRE}$ from competition kinetics (µM) |
|---|---|---|---|
| CREB | 1.2 ± 0.1 | 1.4 ± 0.3 | 1.1 ± 0.1 |
| CREB-FD | 1.11 ± 0.04* | 1.3 ± 0.4 | 1.07 ± 0.03 |
| bZIP | 2.3 ± 0.3 | 0.26 ± 0.08 | 0.50 ± 0.02 |
| bZIP-FD | 2.3 ± 0.08 | 0.36 ± 0.03 | 0.44 ± 0.04 |
| CREBΔQ1 | 1.66 ± 0.06 | 0.75 ± 0.02 | - |
| CREBΔQ2 | 1.36 ± 0.07 | 0.52 ± 0.04 | - |
| CREBΔKID | 1.5 ± 0.1 | - | - |
| CREB-GS-KID | 1.46 ± 0.07 | 0.7 ± 0.1 | - |

**Table S2:** Table of oligonucleotide sequences used in this study. Note, for CRElin and nCRE, only the forward sequence is shown, but are annealed with an oligonucleotide that is an exact reverse complement of the shown sequence. If fluorescently labelled, the fluorophore was added to the 5' most residue.

| Name | Sequence (5'-3') |
|---|---|
| CREh | CCTGACGTCAGCCCCCTGACGTCAGG |
| CRElin | CCTGACGTCATCGG |
| nCRE | GGCTAAAGCATTCT |

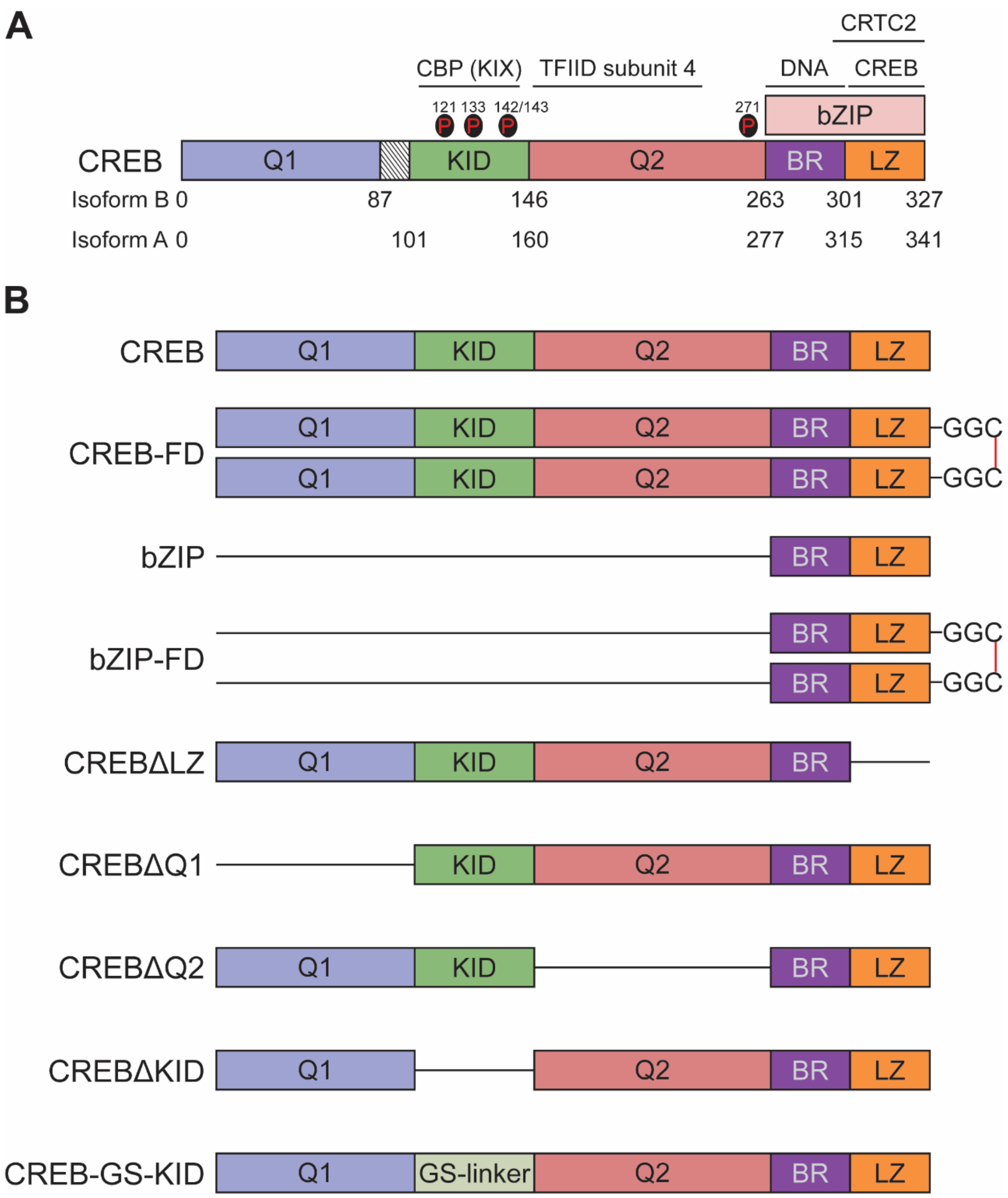


**Figure S1: Overview of CREB constructs. A.** Diagram of CREB domains with length in amino acid numbers, with relevant post translational modification sites and interaction partners. The shaded area between domains Q1 and KID is the isoform A-specific domain. **B.** Diagrams of all mutants used in this study. Deleted domains are replaced with lines. Note that the GGC extension in FD ("forced-dimers") mutants is used to create covalently linked dimers via disulfide bonds (shown in red).

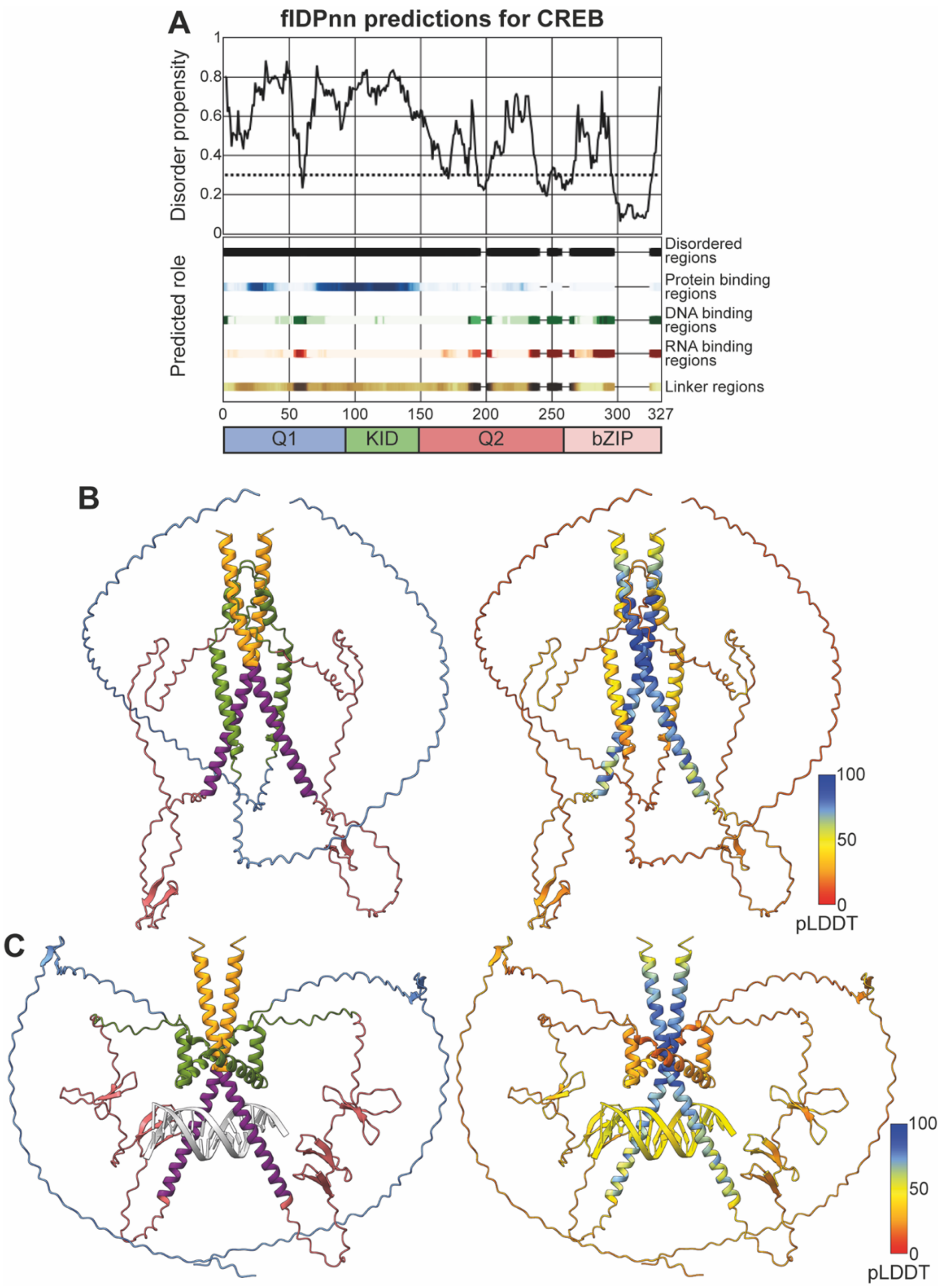


**Figure S2: Predictions of CREB structural properties. A.** Predictions from the fIDPnn (1) webserver show CREB as a mostly disordered protein, but with the leucine zipper helical. Nucleic acid binding regions correctly match the bZIP DNA binding domain and the protein binding regions match the KID activation domain, which interacts with CBP. **B.** AlphaFold3 (2) prediction of the CREB dimer structure, colored according to CREB domains on the left (following the colour scheme in (**A.**)) and coloured according to pLDDT confidence score on the right. As per the fIDPnn prediction, the Q1 and Q2 domains are mostly disordered with some ß -sheets, with the bZIP domain forming a coiled-coil structure. Although confidence score are generally low, this aligns well with previous reports (3,4). **C.** AlphaFold3 prediction of CREB dimers bound to DNA. The bZIP adopts its known coiled-coil domain and the Q1/Q2 domains remain mostly disordered with sparse ß-sheets. Interestingly, the KID domain is predicted to be displaced by DNA compared to (**B.**), although the confidence score is relatively low.

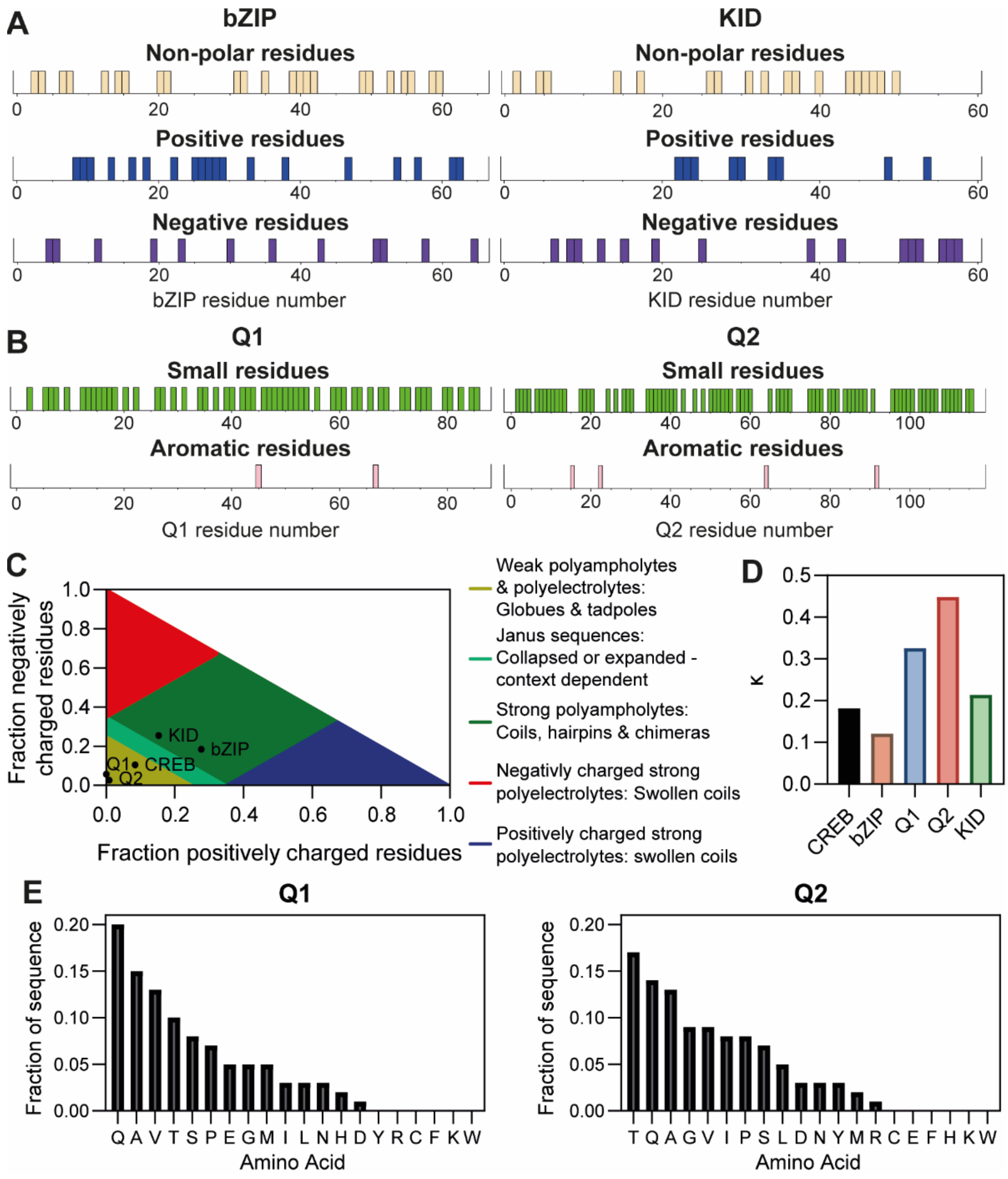


**Figure S3: Overview of CREB domain properties. A.** Repartition of charged and non-polar residues along the bZIP and KID domains, obtained via PEPinfo (5). The former is mostly positive (typical of a DNA binding domain) and the latter mainly negative (typical of an activation domain) but specifically at two patches at the N- and C-termini. **B.** Q1 and Q2 residue repartition obtained via PEPinfo (5). These domains are mainly made of small uncharged residues (see (**C.**) for charge), with the occasional aromatic residue. **C.** CIDER (6) analysis of the CREB sequence. As expected, the bZIP and KID domains are strong polyampholytes due to their numerous charged residues. Overall CREB is considered a weak polyampholyte due to the contribution of Q-rich domains which contain very few charges. **D.** κ values from the CIDER analysis, indicative of the distribution of charges throughout sequences. As indicated by (**A.**), the bZIP domain has more spread-out charges than KID and hence a lower κ value. Q1 and Q2's higher values probably stem more from a low total amount of charges than their distribution. **E.** Fraction of each amino acid content in the two Q-rich domains. As expected, small, uncharged residues dominate (A, G, V, T), and there are very few charged (D, E, K, H, R) or aromatic (H, F, W, Y) residues.

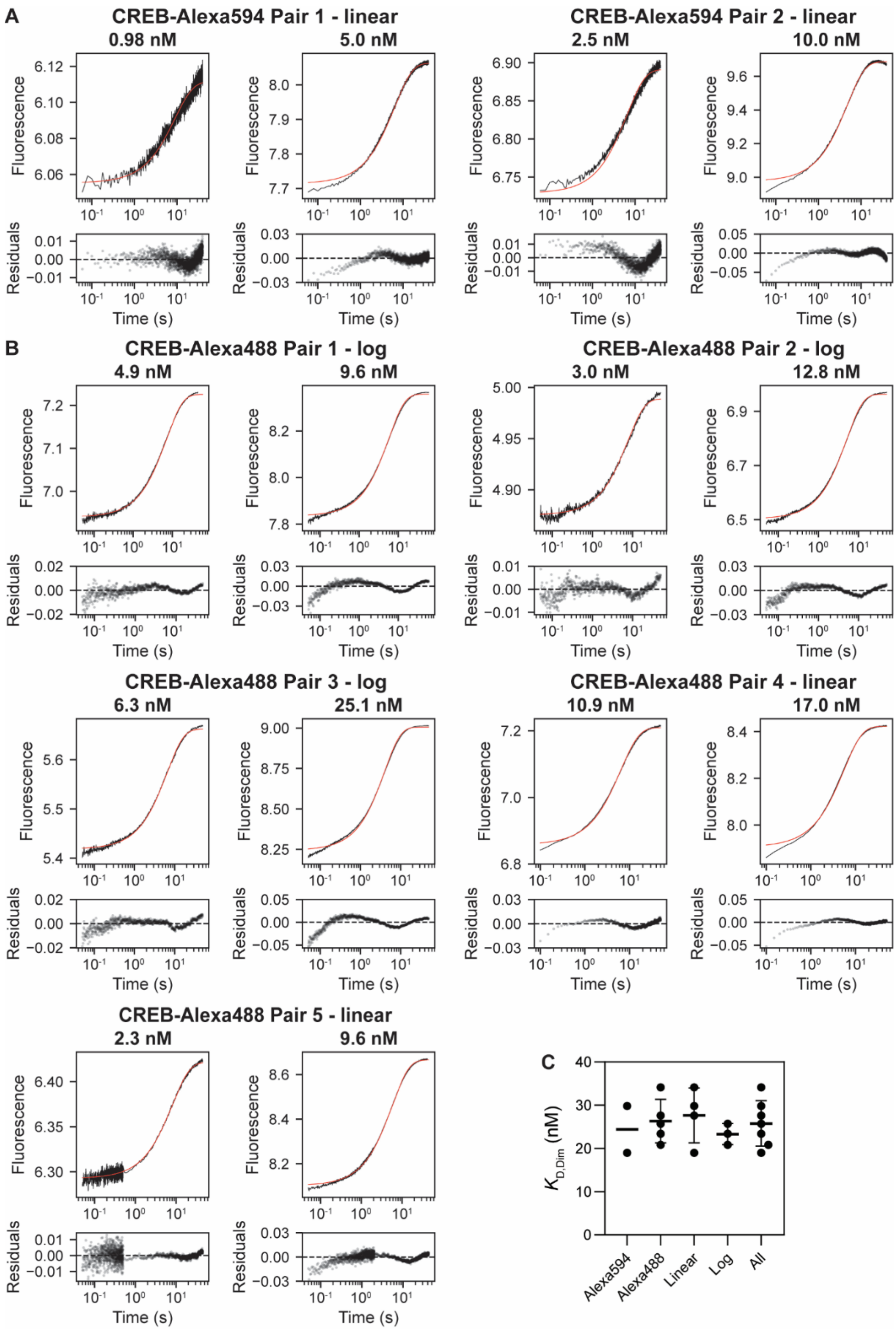


**Figure S4: All pairs of stopped-flow fluorescence relaxation dilution traces for A.** AlexaFluor®594-CREB and **B.** AlexaFluor®488-CREB. Red line indicated the best fit for the pair to the model derived in Eq. 5. Concentrations indicated in the title are those in the optical cell (post-dilution). Associated residuals are displayed below. **C.** $K_{D,dim}$ values derived from the fits in **A.** and **B.**, and are robust to changes in fluorophore and data collection strategy (measurements on a linear or logarithmic time-scale). Individual repeats are shown as datapoints. The average for each grouping is shown as a line, with error bars representing standard deviation, except for AlexaFluor®594-CREB as only two pairs were used.

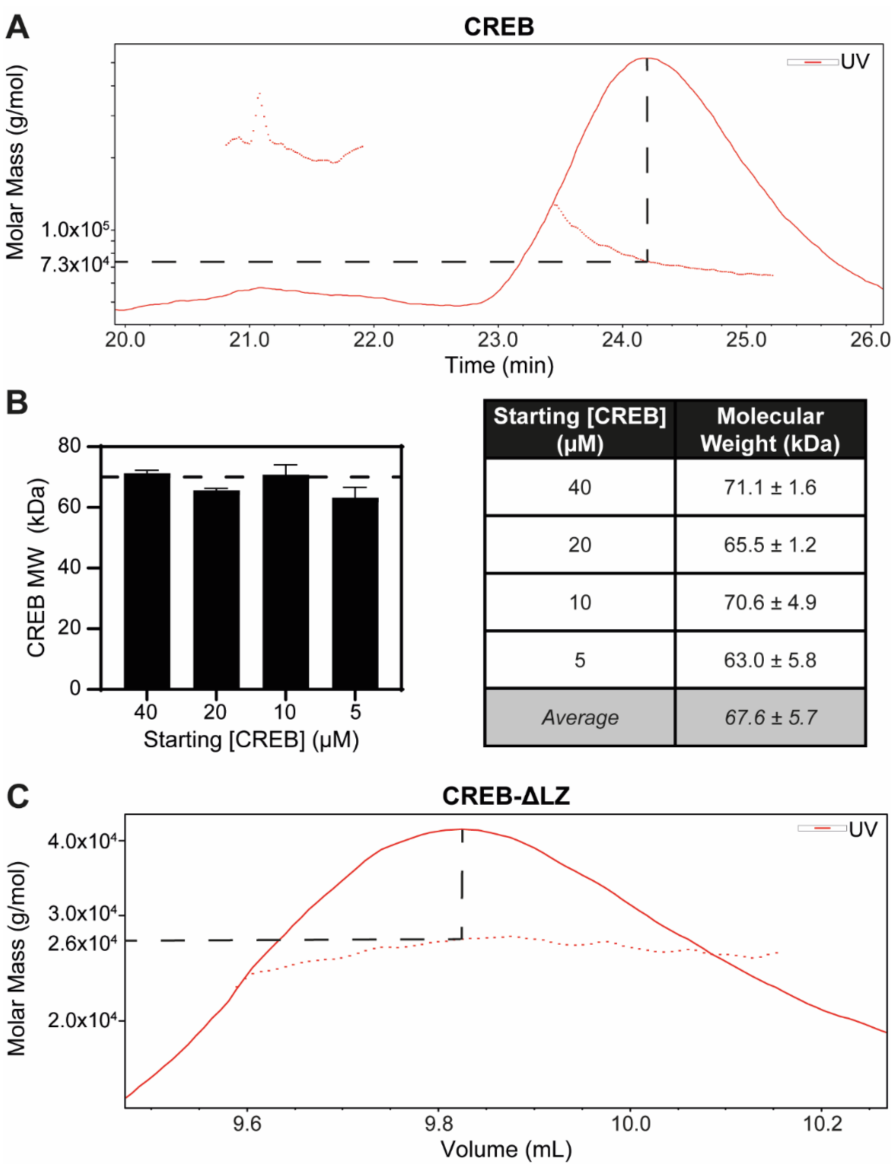


| Starting [CREB] (µM) | Molecular Weight (kDa) |
|---|---|
| 40 | 71.1 ± 1.6 |
| 20 | 65.5 ± 1.2 |
| 10 | 70.6 ± 4.9 |
| 5 | 63.0 ± 5.8 |
| *Average* | *67.6 ± 5.7* |

**Figure S5: SEC-MALS experiments suggest CREB is dimeric in solution.** Protein was loaded onto a Superdex200 Increase 10/300 GL column (GE Healthcare) column at a maximum concentration of 40 µM. Elution with biophysical buffer was monitored via online static light-scattering (DAWN HELEOS-8+, Wyatt Technology), differential refractive index (Optilab T-rEX, Wyatt Technology) and UV (SPD-20A, Shimadzu) detectors. Data were analysed using the ASTRA package v6 (Wyatt Technology). **A.** A SEC-MALS spectrum of CREB at 40 µM shows a large peak at ~73 kDa, roughly corresponding to the dimeric molar mass of ~70 kDa, with a small peak consistent with very minor population of a tetrameric species. **B.** Bar chart of CREB molecular weight from serial dilution experiments, with the expected 70 kDa dimeric weight as a dashed line. Values are also displayed in a table (right). Below 5 µM CREB, no peaks were observed. **C.** SEC-MALS spectrum for CREB-ΔLZ shows a peak at the expected 26 kDa monomeric molar mass.

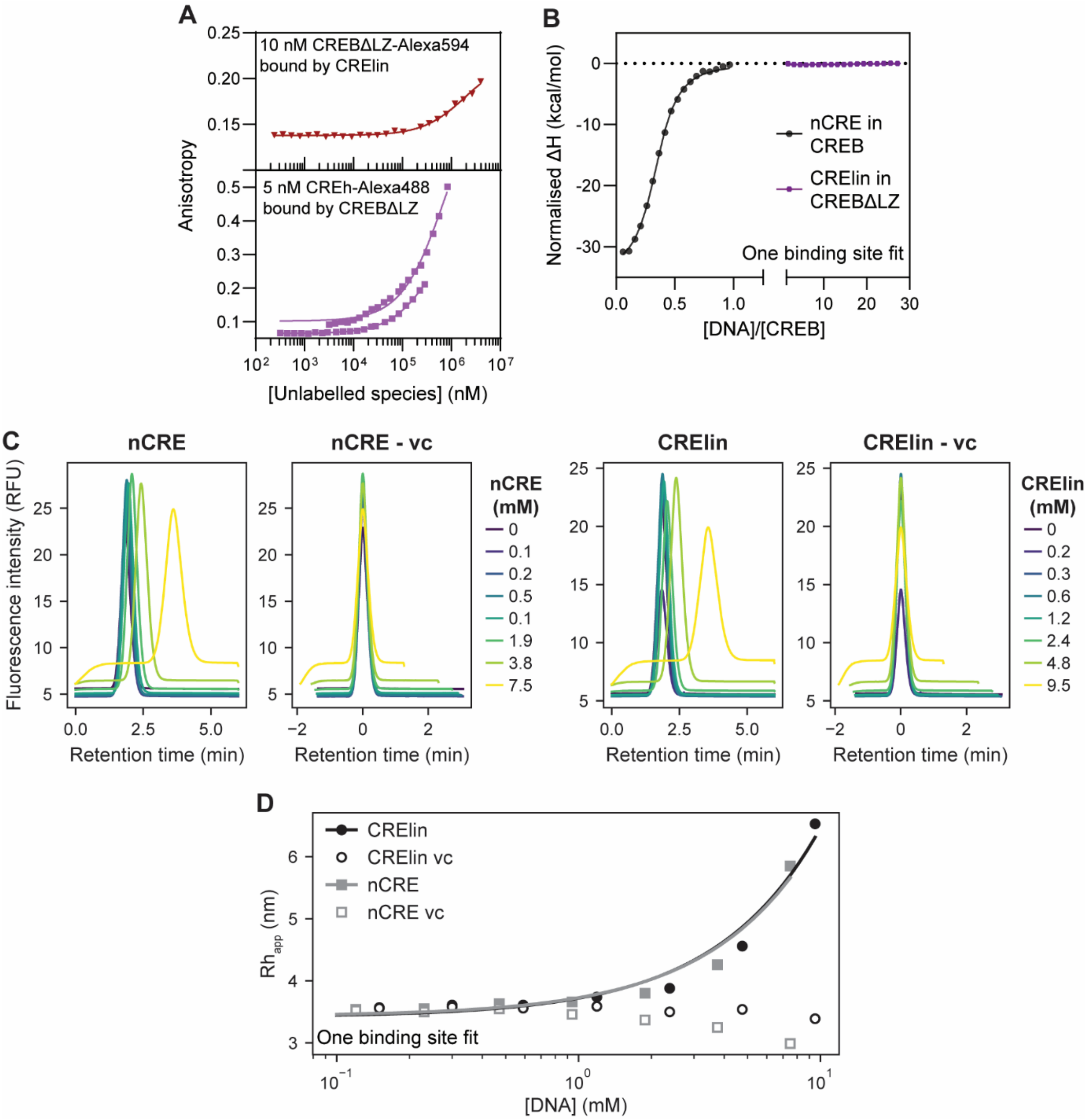


**Figure S6: Lack of evidence for CREBΔLZ binding to DNA. A.** Fluorescence anisotropy equilibrium binding curves, using fluorescently labelled CREBΔLZ binding to CRElin (top) or unlabeled protein binding to fluorescently-labelled CREh (bottom). Anisotropy signals start to increase in the high µM range. An upper saturation plateau is never reached. **B.** ITC titration of CREBΔLZ with CRElin (**right**) shows no heat change. CREB concentrations were above 100 µM, where fluorescence anisotropy is appears increased in **A**. This suggests binding without associated heat change (note heat change is observed titrating CREB with nCRE, **left**), or that anisotropy increases do not result from binding. **C.** Raw FIDA fluorescence signal for nCRE (left 2 panels) and CRElin (right 2 panels), with viscosity corrections (vc). Once viscosity-corrected, there is no change compared to measurements without DNA. **D.** Titrations of Alexa Fluor® 488-labelled CREBΔLZ with unlabeled CRElin and nCRE oligos using Flow-induced dispersion analyses (FIDA). FIDA measures changes in hydrodynamic radius, ($Rh_{,app}$) upon binding. FIDA was performed on a Fida 1 instrument (Fidabio) by the facility at Harwell Research Complex. These titrations show apparent increases in $Rh_{,app}$ at mM DNA concentrations (solid datapoints) that disappear once solution viscosity is considered (vc, open datapoints).

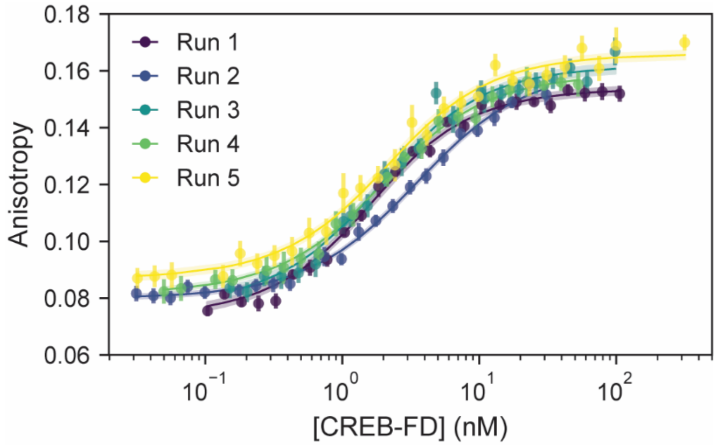


**Figure S7**: **Fluorescence anisotropy equilibrium curves for CREB-FD binding to Alexa Fluor 488-labelled CREh.** Lines represent fit to a single binding site model (Eq. 3). These show high affinity for target DNA, in the low nM range.

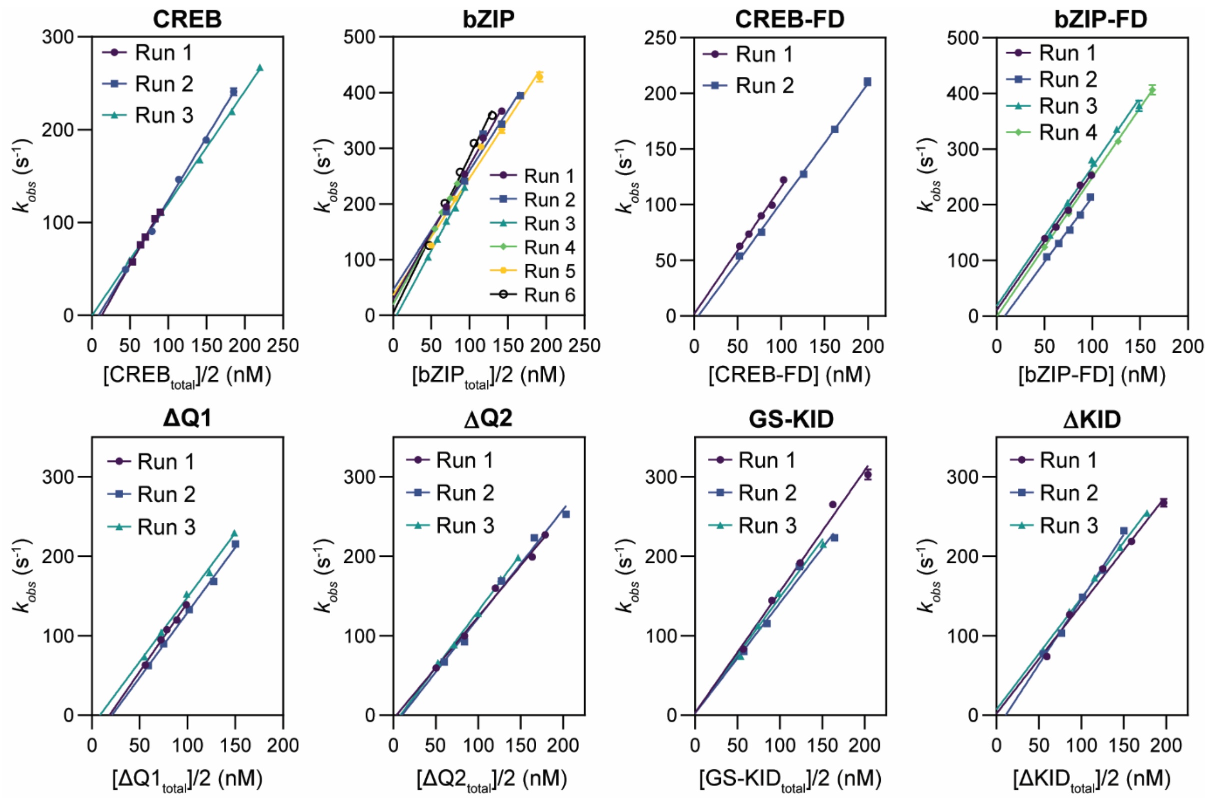


**Figure S8: Observed association rates for CREB pseudo wild-type and mutants to 5 nM Alexa Fluor® 488-labelled CREh.** Observed rate constants ($k_{obs}$), obtained from single exponential decay fits to stopped-flow kinetic traces, show a linear dependence on protein concentration. Results of independent experiments performed on different days (with different protein batches) are each fit to a straight line.

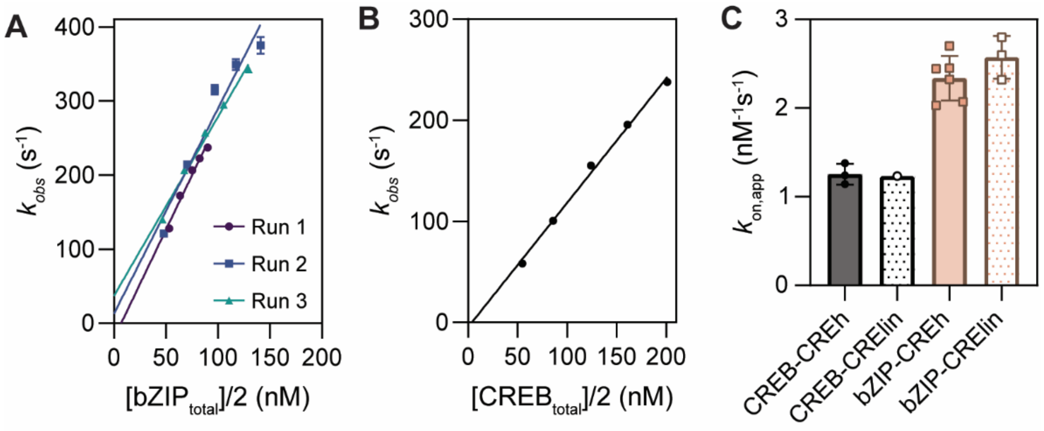


**Figure S9: Association rate constant is similar for hairpin and linear DNA.** Extracted observed rate constants ($k_{obs}$), both for **A.** CREB-bZIP and **B.** CREB with 5 nM Alexa Fluor® 488-labelled CRElin show a linear dependence on concentration, where the gradient represents $k_{on}$. **C.** Bar chart showing $k_{on}$ averages from the individual datapoints for CREB and bZIP. Binding to labelled CREh is shown as full bars, and binding to CRElin is shown as spotted bars. All errors are standard deviation from replicate averages, except for CREB binding to CRElin due to the single experiment instance.

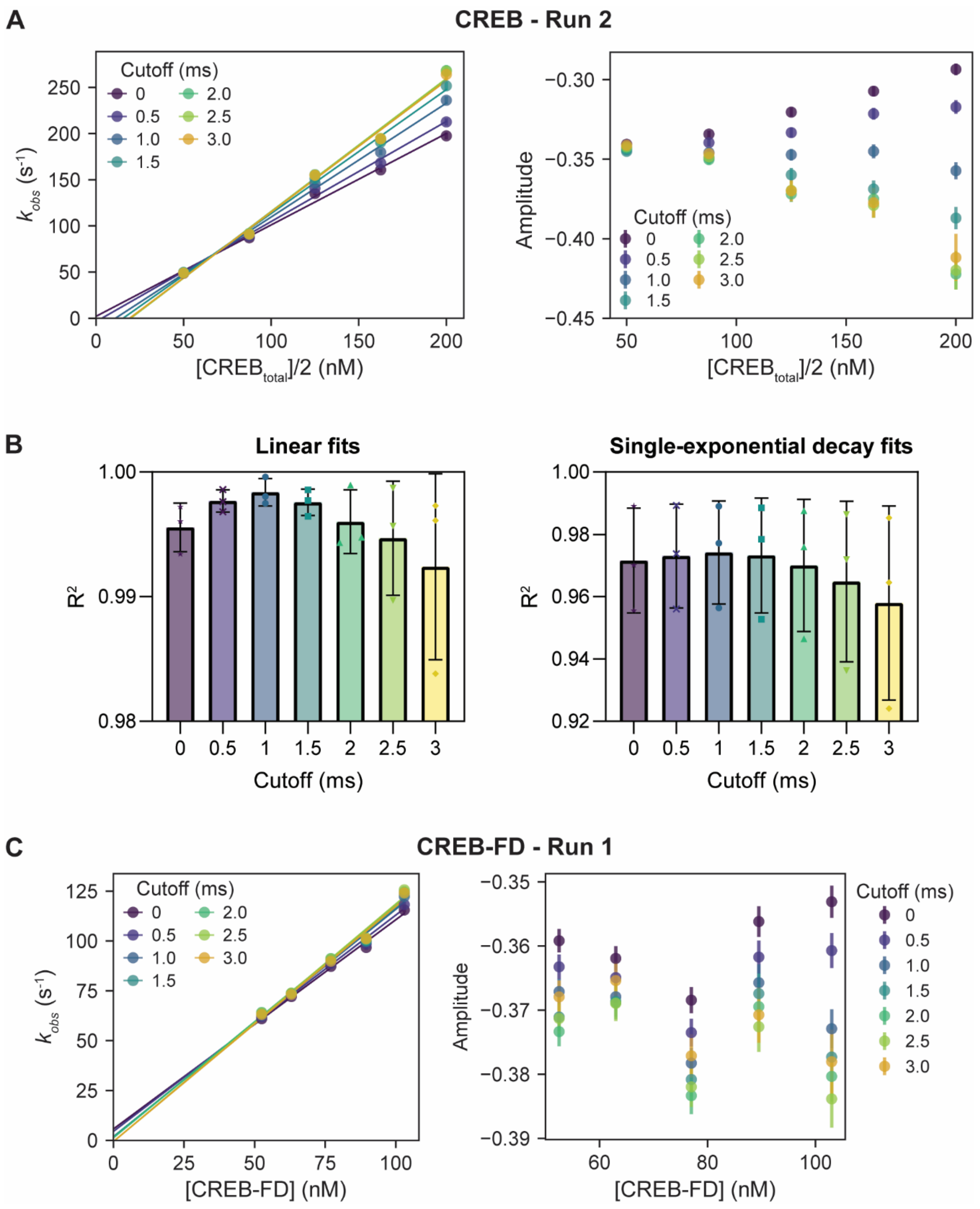


**Figure S10: Determining optimal cutoff for association kinetic fitting. A.** $k_{obs}$ (left) and amplitude (right) of single exponential decay fits of CREB-CREh association traces, when using different cutoff times to account for apparatus mixing time. Decreasing cutoff time decreases $k_{obs}$ and shifts the y-intercept closer to 0 (left). The amplitude, which is expected to be roughly constant at these concentrations due to saturated binding, is most consistent with 1 ms cutoff. **B.** Fit quality assessments ($R^2$ values) for the linear concentration dependence (left) and exponential decay observed in kinetic traces (right) of CREB. All values are similar and within error, but the best fit is consistently a 1 ms cutoff. Bars are the averages of the plotted replicates, and errors display standard deviation. **C.** Same as (**A.**) but with CREB-FD. The cutoff dependence is reduced.

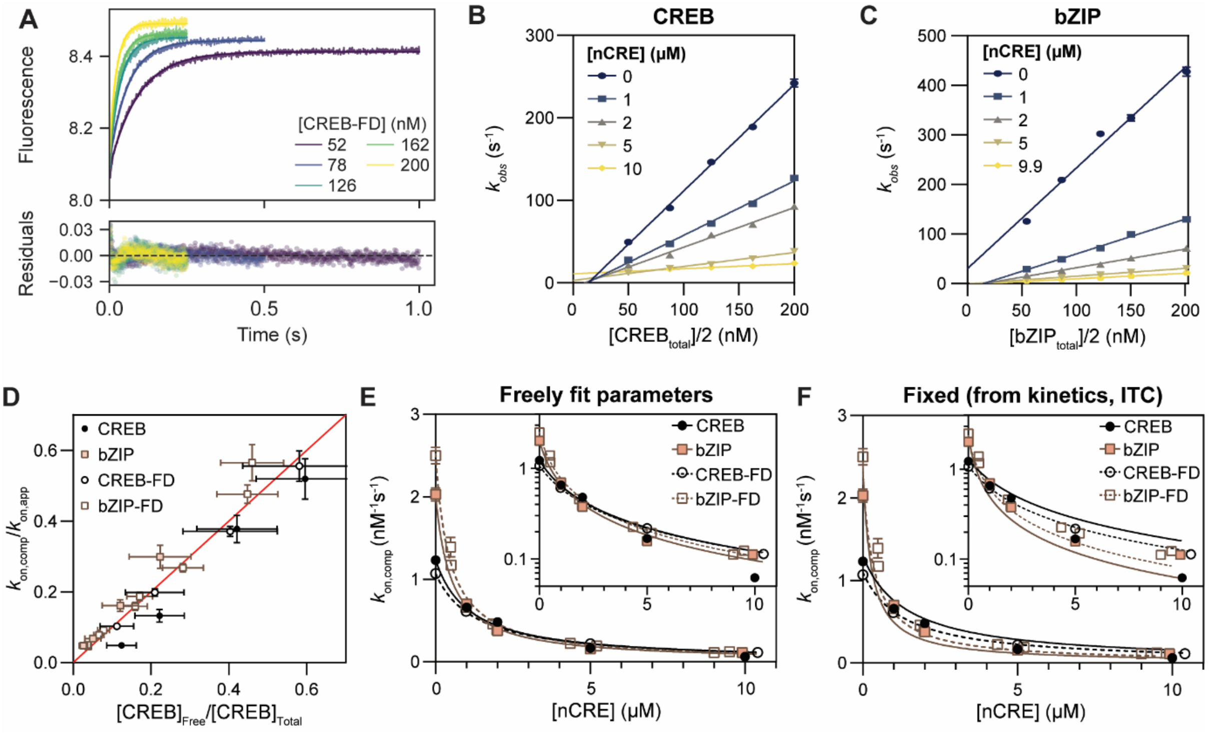


**Figure S11: Mimicking the target-search of CREB *in-vitro*. A.** Association traces of CREB-FD to 5 nM AlexaFluor®488-CREh in the presence of 5 µM unlabeled nCRE competitor. Concentrations are those in the optical cell. Lines show fit to a single exponential decay function. Residuals are displayed in the lower panel. **B and C.** $k_{obs}$ from association experiments at various [nCRE] plotted against protein concentration for CREB (**B.**) and CREB-bZIP (**C.**) demonstrate a linear dependence on protein concentration, with the gradient decreasing with increasing nCRE concentrations. **D.** Ratio of inhibited against uninhibited $k_{on}$ against the expected fraction of unbound CREB (calculated from $K_{D,nCRE}$). Errors were propagated according to standard rules. Data are consistent with the red line, y=x. **E.** and **F.** Decrease of association rate constants to CREh with increasing nCRE competitor ($k_{on,comp}$) for CREB, bZIP and respective FD mutants. Data is fitted to the model derived in Eq. 4. **E.** The extent of inhibition is comparable for forced dimers and pseudo wild-type. **F.** Data broadly match predictions based on independent estimates of $K_{D.nCRE}$ and $k_{on}$ from ITC and kinetic experiments.

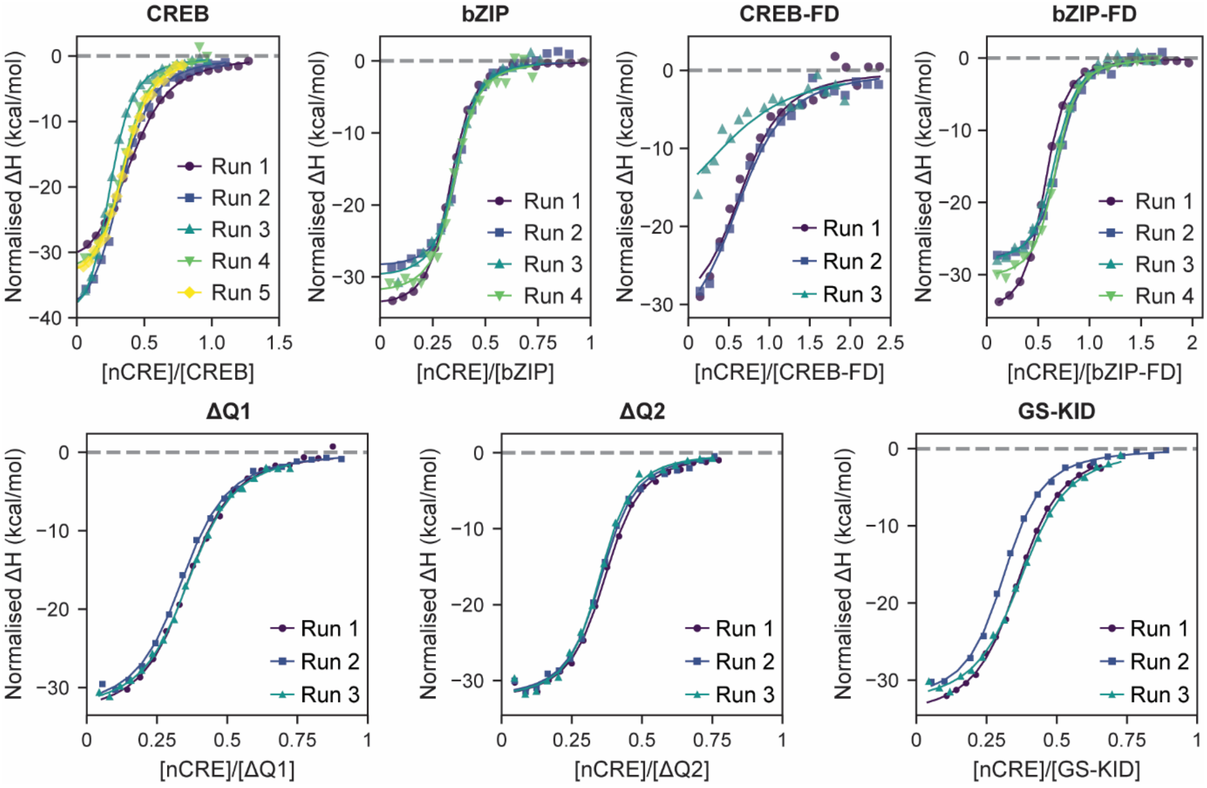


**Figure S12: ITC titrations for CREB mutants.** Normalised ΔH in kcal/mol is plotted against the titrant [nCRE] to titrand [protein] ratio, for each mutant. Lines represent fit to a one binding site model.

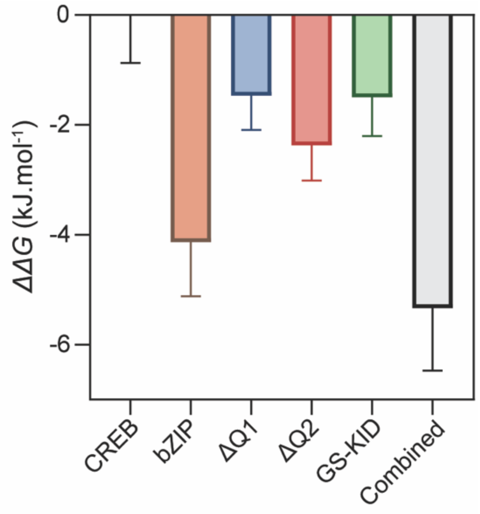


**Figure S13: ΔΔG values of binding to nCRE for domain mutants**. Comparison is to CREB. The sum of ΔΔG for each mutant slightly exceeds that of the bZIP, but is within error.

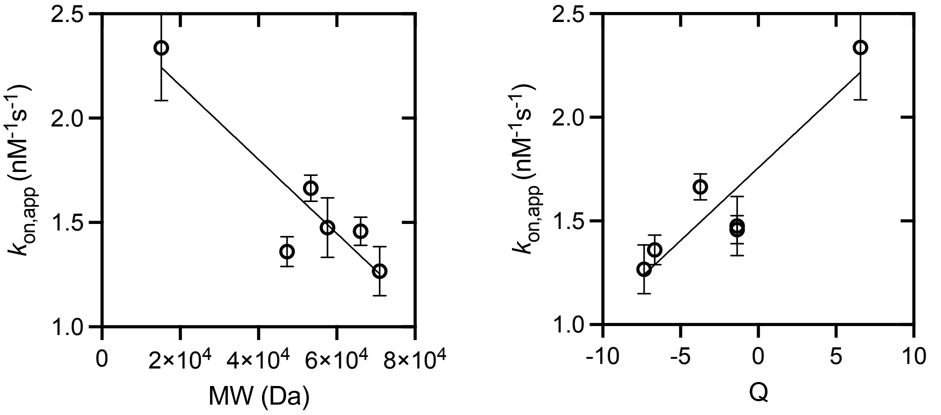


**Figure S14: Association rate constants with target DNA correlate with both molecular weight (left) and net charge of the protein (right).** Correlations are statistically significant in both cases ($p = 0.01$, $n = 6$, two-tailed).

### Equation derivations

#### $k_{on}$ inhibition of binding to target DNA by non-target DNA

To derive the model used to fit the $k_{\mathrm{on,comp}}$ of association to labelled CREh in the presence of non-target competitor nCRE, we make the assumption that the $k_{\mathrm{on,comp}}$ would be directly proportional to the amount of free CREB non-sequestered by nCRE as in Eq. S1:

$$k_{\mathrm{on,comp}} = k_{\mathrm{on}} \times \frac{[CREB]_{free}}{[CREB]_{total}} \qquad (S1)$$

The equilibrium dissociation constant for CREB binding to nCRE is equal to:

$$K_{\mathrm{D,nCRE}} = \frac{[CREB]_{free} \times [nCRE]_{free}}{[CREB{:}nCRE]} \qquad (S2)$$

Conservation of CREB molecules requires that

$[CREB{:}nCRE] = [CREB]_{total} - [CREB]_{free}$ $(S3)$,

Substitution and rearrangement leads to Eq. S4

$$\frac{[CREB]_{total}}{[CREB]_{free}} = \frac{[nCRE]_{free} + K_{\mathrm{D,nCRE}}}{K_{\mathrm{D,nCRE}}} \qquad (S4)$$

Since the nCRE concentration remains almost constant, Equation S4 can be substituted into Eq. S1 to yield:

$$k_{\mathrm{on,comp}} = k_{\mathrm{on,app}} \times \frac{K_{\mathrm{D,nCRE}}}{[nCRE] + K_{\mathrm{D,nCRE}}} \qquad (S5)$$
